# Adaptation to skin mycobiota promotes antibiotic tolerance in *Staphylococcus aureus*

**DOI:** 10.1101/2024.05.03.592489

**Authors:** Caitlin H. Kowalski, Susannah Lawhorn, T. Jarrod Smith, Rebecca M. Corrigan, Matthew F. Barber

## Abstract

The microbiota can promote host health by inhibiting pathogen colonization, yet how host-resident fungi, or the mycobiota, contribute to this process remains unclear. The human skin mycobiota is uniquely stable compared to other body sites and dominated by yeasts of the genus *Malassezia*. We observe that colonization of human skin by *Malassezia sympodialis* significantly reduces subsequent colonization by the prominent bacterial pathogen *Staphylococcus aureus*. *M. sympodialis* secreted products possess potent bactericidal activity against *S. aureus* and are sufficient to impair *S. aureus* skin colonization. This bactericidal activity requires an acidic environment and is exacerbated by free fatty acids, demonstrating a unique synergy with host-derived epidermal defenses. Leveraging experimental evolution to pinpoint mechanisms of *S. aureus* adaptation in response to the skin mycobiota, we identified multiple mutations in the stringent response regulator Rel that promote survival against *M. sympodialis*. Similar Rel alleles have been reported in *S. aureus* clinical isolates, and natural Rel variants are sufficient for tolerance to *M. sympodialis* antagonism. Partial stringent response activation underlies tolerance to clinical antibiotics, with both laboratory-evolved and natural Rel variants conferring multidrug tolerance. These findings demonstrate the ability of the mycobiota to mediate pathogen colonization resistance, identify new mechanisms of bacterial adaptation in response to fungal antagonism, and reveal the potential for microbiota-driven evolution to shape pathogen antibiotic susceptibility.

**Highlights:**

- *M. sympodialis* reduces colonization of human skin by *S. aureus*
- Bactericidal activity of *M. sympodialis* is exacerbated by features of the skin niche
- *S. aureus* Rel variants are sufficient for tolerance to *Malassezia* antagonism
- Evolved tolerance to yeast antagonism coincides with *S. aureus* multidrug tolerance

## Introduction

*Staphylococcus aureus* is the primary cause of human skin and soft tissue infections, resulting in approximately 500,000 hospitalizations annually in the United States ^1^, and in 2019 was associated with more than one million deaths globally ^2^. Despite the ability of *S. aureus* to thrive within wounds and to asymptomatically colonize the nares, colonization of healthy skin typically occurs transiently ^3, 4^. Several features of the healthy epidermal environment are predicted to contribute to this colonization deficiency, including acidic pH, antimicrobial fatty acids, and host-produced antimicrobial peptides ^5–7^. In addition to these innate host features, some skin-resident bacteria possess the ability to inhibit pathogen colonization, a process referred to as colonization resistance ^8^. Microbial factors can act by inhibiting *S. aureus* directly ^9, 10^, disrupting biofilm formation ^11^, activating host defenses ^12^, or modulating *S. aureus* virulence ^13, 14^. Indeed, the application of a naturally antagonistic skin-resident bacteria is currently in clinical trials for the treatment of *S. aureus* colonization in atopic dermatitis patients ^15^.

Despite several examples of skin-resident bacteria that mediate colonization resistance against *S. aureus* ^8^, and the critical need to identify novel antimicrobials ^16^, relatively little is known about how skin-resident fungi contribute to host health through interactions with invading pathogens. While fungi have historically been overlooked due to their low abundance compared to resident bacteria, human skin contains a comparatively larger proportion of fungi relative to other body sites ^17, 18^. Notably, this skin mycobiota is largely dominated by a single genus of yeast, *Malassezia* ^19, 20^. Ubiquitous colonizers of mammalian skin, *Malassezia* yeasts are well-adapted to the dermal environment. Firstly, *Malassezia* spp. have some of the smallest genomes of known free-living fungi, a feature commonly observed among microorganisms that adapt to a specific host environment ^21, 22^. Second, *Malassezia spp.* have horizontally acquired genetic material from skin-resident bacteria, which may further contribute to niche-specific adaptation ^21, 23^. Finally, *Malassezia* have lost the ability to produce fatty acids *de novo* and are dependent on exogenous lipid sources, such as those abundant on the skin surface ^21, 24^. As these yeasts likely adapted to the skin environment in the presence of other resident microbes, we hypothesized *Malassezia* have evolved mechanisms for intermicrobial competition. Consistent with this hypothesis, one of the most prevalent human skin colonizers, *Malassezia globosa*, was reported to secrete a protease capable of disrupting *S. aureus* biofilms ^25^, and recent studies suggest a possible protective role for *Malassezia* against colonization by the emerging fungal pathogen *Candida auris* ^26, 27^. Beyond these examples, it is unknown if *Malassezia* produce other antimicrobial effectors capable of inhibiting bacterial colonization or how pathogens may adapt to antagonists within the microbiota.

Here, we use living human skin biopsies from healthy donors and *in vitro* host-like conditions to investigate the ability of *Malassezia* to inhibit *S. aureus* colonization and growth. We further investigate the potential for microbiota antagonism to shape pathogen evolution by selecting for tolerance in *S. aureus* exposed to *Malassezia* or its secreted products. We report that the presence of *M. sympodialis* in a mixed species biofilm with *S. aureus in vitro*, or co-colonized with *S. aureus* on the epidermis, results in significantly reduced *S. aureus* growth and colonization, respectively. *M. sympodialis* secretes potent antimicrobials *in vitro* that are uniquely active against *S. aureus* within the physical constraints of the dermal environment and significantly reduce *S. aureus* skin colonization. Long-term exposure to *M. sympodialis* or repeated exposure to its secreted antimicrobial products selects for stable tolerance in *S. aureus* through partial activation of the stringent response brought about by mutations in the GTP pyrophosphokinase Rel. Alleles like those identified in our experimentally evolved tolerant strains have been identified in *S. aureus* clinical isolates and both are sufficient for stringent response activation, tolerance to *M. sympodialis,* and multidrug tolerance. This work proposes that skin resident fungi contribute to colonization resistance through production of potent antimicrobials, highlighting the important role that often-overlooked fungi play at the interface of the microbiota and host health. Furthermore, our findings illustrate the potential for intermicrobial antagonism to shape pathogen evolution and antibiotic susceptibility.

## Results

### Malassezia sympodialis inhibits Staphylococcus aureus growth in vitro

To test the hypothesis that *Malassezia* species can alter *S. aureus* growth, we first selected a clinical isolate of *S. aureus,* NRS193, as our wild type (WT) strain (**Table S1**). This community-associated methicillin resistant *S. aureus* (CA-MRSA) strain is a USA400 isolate from a pneumonia patient and was selected because it is closely related to the reference strain MW2, is not laboratory adapted, and did not originate from a skin infection ^28^. We inoculated NRS193 adjacent to 72-h colonies of *Malassezia sympodialis*, *Malassezia furfur*, or *Malassezia pachydermatis* grown on mDixon agar. This media is typically used to culture fastidious *Malassezia* and recapitulates the acidic and lipid-rich environment of the skin. After 24-h of *S. aureus* growth, only *M. sympodialis*, and not *M. furfur* or *M. pachydermatis*, inhibited adjacent growth of *S. aureus* (**Fig. 1A**). *M. sympodialis* is one of the three most prevalent species of *Malassezia* on healthy human skin, while *M. furfur* is less abundant but more easily culturable ^21^. In contrast*, M. pachydermatis* is a common colonizer of other mammals such as canines (**Table S1**) ^29^. Notably, *M. sympodialis* did not impact the adjacent growth of *Escherichia coli* or the abundant skin commensal *Staphylococcus hominis* (**Fig. 1B**). These data indicate that established colonies of *M. sympodialis* can reduce the growth of adjacent *S. aureus*.

**Figure 1.**
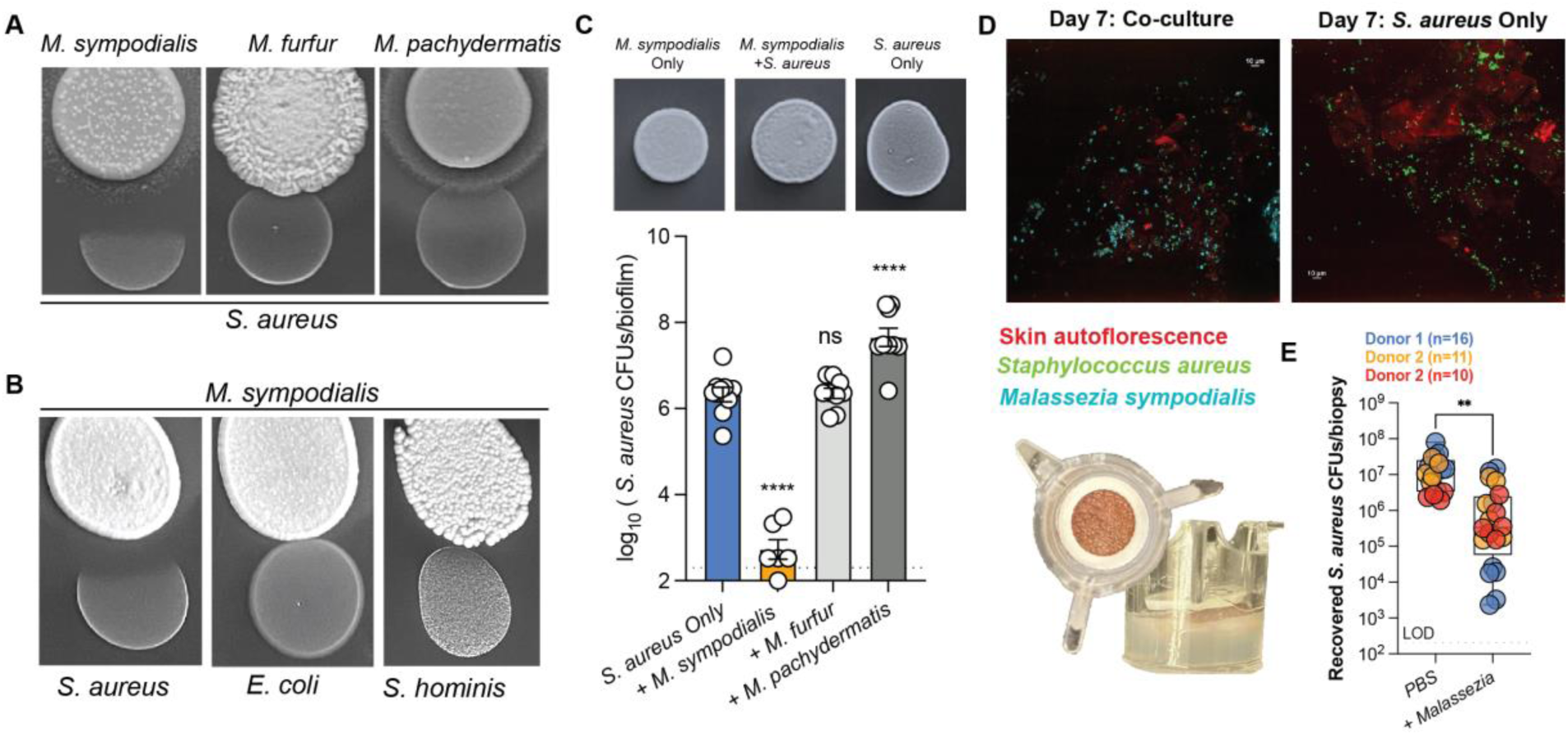
*Malassezia sympodialis* inhibits the pathogen *S. aureus in vitro* and on human skin. **A.** Representative images of *S. aureus* NRS193 24-h growth adjacent to *M. sympodialis, M. furfur, or M. pachydermatis* colonies that had been grown on mDixon agar for 72-h before *S. aureus* inoculation. **B.** Representative images of *S. aureus* NRS193, *E. coli*, Nissle or *S. hominis* SK119 24-h growth adjacent to *M. sympodialis* that had been grown on mDixon agar for 72-h prior to bacterial inoculation. **C.** Top: representative images of 6-d colony biofilms on mDixon agar. Bottom: *S. aureus* NRS193 CFUs enumerated after 6-d growth in a colony biofilm alone (*S. aureus only*) or with *M. sympodialis, M. furfur,* or *M. pachydermatis*. One-way ANOVA with Dunnett’s multiple comparisons test. ns: not significant, ****: p<0.0001. Data pooled from three separate experiments where each point represents one colony biofilm. (n=9 per condition). Four biofilms with *M. sympodialis* had no recovered *S. aureus* above the limit of detection (200 CFUs/mL). **D.** Representative maximum projections images of the skin surface (red) at day 7 colonized with *M. sympodialis* (cyan) and/or *S. aureus* (green). Scale bar is 10 µm. **E**. Left: image of Nativeskin Access (Genoskin) human skin biopsy, Right: Recovered *S. aureus* CFUs per human skin biopsy with data combined from three experiments with three skin donors. Each point represents a skin biopsy. (PBS n=17, +*Malassezia* n=20). Colors correspond to respective donor. **:p<0.01 calculated with unpaired, parametric t-test. Whiskers are Min to Max with all points shown. LOD: limit of detection (200 CFUs/mL)

To determine if *Malassezia* impacts *S. aureus* growth when inoculated simultaneously, we established mixed colony biofilms on mDixon agar of *S. aureus* NRS193 with *M. sympodialis, M. furfur*, or *M. pachydermatis*. The presence of *M. sympodialis* significantly reduced the recovery of *S. aureus* (measured as colony forming units (CFUs)/ biofilm) compared to *S. aureus* alone after 6 days (**Fig. 1C**) (n=9 per group, with no *S. aureus* recovered above the limit of detection in four biofilms with *M. sympodialis*). In contrast, co-culture with *M. furfur* or *M. pachydermatis* did not significantly reduce the recovery of *S. aureus* relative to its growth in monoculture. In fact, culturing with *M. pachydermatis* appeared to increase the recovery of *S. aureus* (**Fig. 1C**). While mDixon is the classic media for culturing *Malassezia*, we also performed mixed colony biofilm assays on a modified potato dextrose agar (mPDA), previously described ^30^. We observed similar results, with a reduction in *S. aureus* recovery with *M. sympodialis* compared to *S. aureus* alone (**Fig. S1A**). Collectively, we conclude that *M. sympodialis* can antagonize *S. aureus in vitro*.

### Malassezia sympodialis inhibits Staphylococcus aureus colonization of human skin

Based on our *in vitro* observations, we hypothesized that *M. sympodialis* colonization of human skin would reduce subsequent colonization by *S. aureus.* Human skin explants, or biopsies, have previously been used to study host-microbe interactions, including host-*Malassezia* interactions ^31–33^. These explants are viable, full thickness human skin that contain skin appendages, such as follicles, and tissue-resident leukocytes ^32^. Sterilization of skin biopsies (NativeSkin Access, GenoSkin) prior to surgery appears to abolish much of the existing microbiome, as plating on non-selective media did not result in growth of aerobic bacteria or yeast. We inoculated biopsies, prepared from abdominal skin of three female donors, with *M. sympodialis* (n=20 biopsies total from 3 donors) or the vehicle control PBS (n=17 biopsies from 3 donors). *M. sympodialis* was labeled with the cell wall stain calcofluor white at a subinhibitory concentration to confirm colonization on the skin surface through microscopy (**Fig. 1D, Fig. S2A, B**). Six days after inoculation with *M. sympodialis* or PBS, all biopsies were inoculated with *S. aureus* NRS193 engineered to constitutively express green fluorescent protein (GFP). After 24-h of *S. aureus* colonization, biopsies were physically disrupted to remove *S. aureus*. Biopsies pre-colonized with *M. sympodialis* yielded significantly less recovered *S. aureus* (CFUs/mL biopsy) compared to those that received the PBS control (**Fig. 1E**, **Fig. S2C**) (**: p<0.01). These findings suggest, consistent with our *in vitro* observations, that *M. sympodialis* is capable of antagonizing *S. aureus* on the skin surface.

### M. sympodialis secretes potent bactericidal products active against S. aureus

Several skin resident bacteria are known to secrete antimicrobial compounds capable of inhibiting *S. aureus* growth ^9, 10, 15, 34^ and reducing its virulence ^13, 14^. Initial *in vitro* experiments further suggested that the antimicrobial activity of *M. sympodialis* is secreted (**Fig. 1A**). To confirm whether *M. sympodialis* antagonizes *S. aureus* through a secreted factor, we grew *M. sympodialis* in monoculture and collected the cell-free supernatant (CFS) after 96-h. Compared to pH-matched media controls, *M. sympodialis* CFS (Ms-CFS) from mDixon shows dose-dependent antimicrobial activity against *S. aureus* NRS193 after only 2-h treatment (**Fig. 2A**), with ∼ 1000-fold reduction in viable *S. aureus* CFUs in the presence of 50% Ms-CFS. When *M. sympodialis* is grown in modified potato dextrose broth (mPDB) or synthetic sebum media, the Ms-CFS treatment takes 24-h to reduce *S. aureus* NRS193 CFUs by 1000-fold relative to the pH-matched media control (**Fig. S1B, C**). This suggests components in the media can alter yeast production of the antimicrobial or exacerbate the antimicrobial effect contributing to its *in vitro* potency.

**Figure 2.**
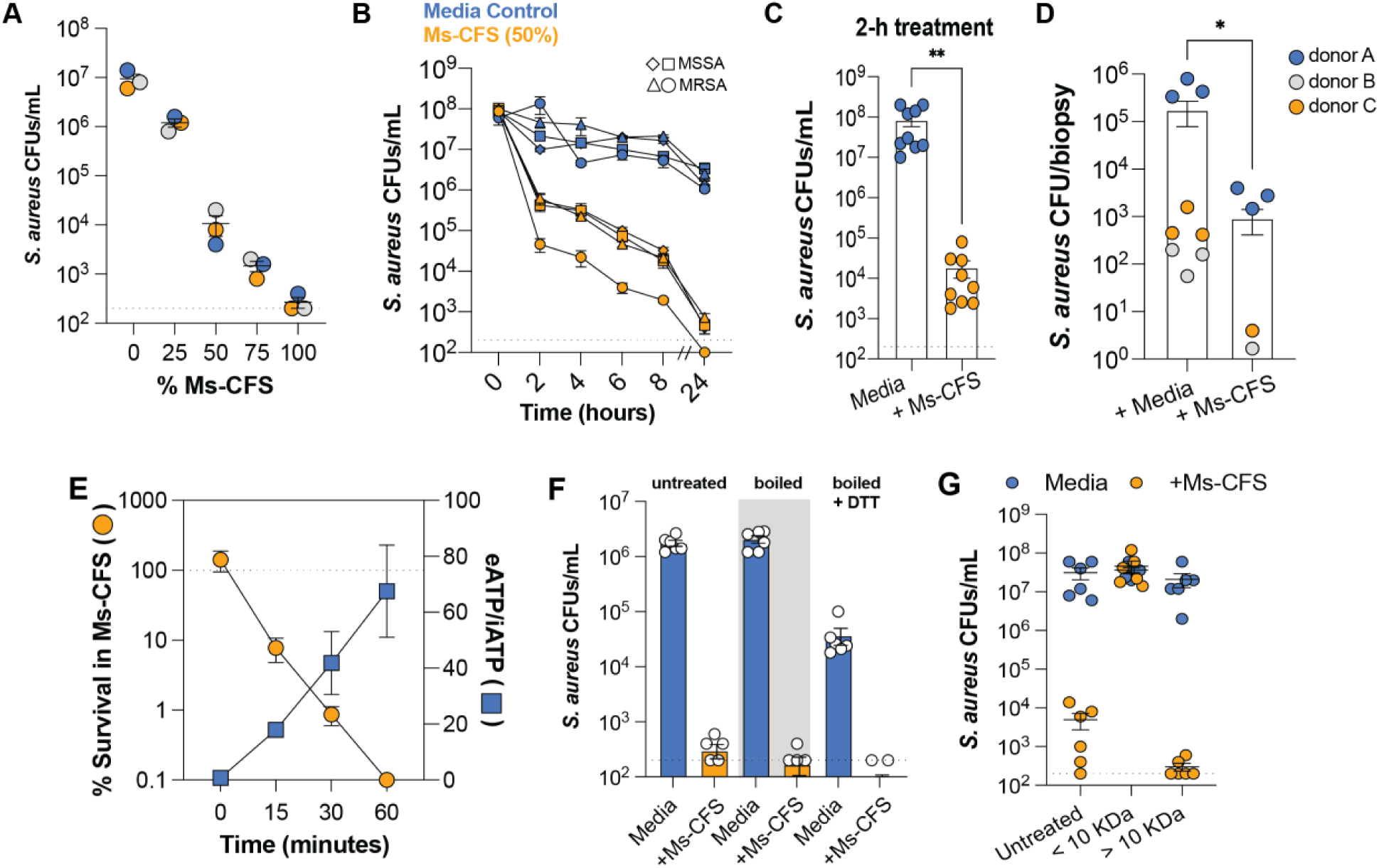
*M. sympodialis* secretes bactericidal products with potent membrane perturbing activity against *S. aureus*. **A.** Cell-free supernatant collected from *M. sympodialis* 96-h monoculture is bactericidal to *S. aureus* NRS193 in a dose-responsive manner (pH 5.5) (n=3). **B.** *S. aureus* CFUs from four separate strains (2 MSSA-diamond, square, 2 MRSA-triangle, circle) quantified over 24-h in the presence of 50% *M. sympodialis* CFS (orange) or pH-matched media control (blue). **C**. *S. aureus* NRS193 (circle strain from C B?) CFUs quantified after 2-h exposure to 50% *M. sympodialis* CFS (orange) or pH-matched media control (blue). **D**. *S. aureus* CFUs (RFP-expressing strain) recovered from the surface of human skin biopsies after 24-h colonization followed by 24-h treatment with either 50% CFS or Media. Each point represents a biopsy colored by donor. n=9 biopsies per group with n=3 biopsies from each donor, 4 biopsies had no recoverable *S. aureus* in the + CFS group and are not shown. **E.** Quantification of *S. aureus* NRS193 CFUs (orange circles) and extracellular (eATP) to intracellular ATP (iATP) ratio (blue squares) over 60 minutes of exposure to 50% *M. sympodialis* CFS. Data is representative of three separate experiments. **F.** *S. aureus* NRS193 CFUs after 2-h exposure to 50% CFS that was untreated, boiled for 15 min. at 95C, or boiled for 15 min. at 95C with 1 mM DTT. Data from three separate treatments of media or CFS. **G.** *S. aureus* NRS193 CFUs after 2-h exposure to 50% CFS that was untreated or separated across a 10 kDa filter with volume adjusted. Data pooled from two experiments, n=6. Dashed line is limit of detection: 200 CFUs/mL, except in E where dashed line denotes 100% survival in CFS. Unpaired, parametric t-tests performed. **: p<0.01, *: p<0.05.

We opted to continue with CFS prepared from *M. sympodialis* grown in mDixon and monitored the antibacterial activity of 50% Ms-CFS treatment over 24-h with four different *S. aureus* strains. With two methicillin sensitive and two methicillin resistant *S. aureus* strains (MSSA and MRSA, respectively), we observed rapid killing during the first 2-h of 50% Ms-CFS treatment (100-to-1000-fold reduction) followed by a more gradual reduction in viability from 2 to 24-h (**Fig. 2B**). Notably, there is no observed growth of any of the *S. aureus* strains in the pH-matched media control (pH 5.5) over 24-h, despite *S. aureus* growing in mDixon at pH 6 (**Fig. S3A**). Given that we observed reproducible rapid killing in the first 2-h of treatment, we decided to use 2-h treatment with 50% Ms-CFS (now referred to as ‘+ Ms-CFS’ unless otherwise noted) for future experiments (**Fig. 2C**). We next sought to determine whether Ms-CFS is sufficient to reduce *S. aureus* colonization on human skin. To this end, we inoculated human skin biopsies with *S. aureus* for 24-h before treating with either the pH-matched media control or Ms-CFS for an additional 24-h. Biopsies were disrupted to recover *S. aureus* from the skin surface, and we observed a significant reduction in *S. aureus* recovered from biopsies treated with the Ms-CFS compared to the media control (**Fig 2D**) (*: p<0.05, n=9 per group from 3 total donors). Of note, the skin colonization experiment was performed with a strain of *S. aureus* expressing red fluorescent protein (RFP) observed to also have *in vitro* susceptibility to *M. sympodialis* (**Fig. S3B**). Together these data indicate that *M. sympodialis* cultured *in vitro* secretes antimicrobials with bactericidal activity against *S. aureus* that are capable of reducing colonization on human skin.

### M. sympodialis antimicrobial activity results in S. aureus membrane perturbation

The rapid killing of *S. aureus* by *M. sympodialis* CFS appears to be independent of bacterial growth, as *S. aureus* CFUs do not increase within the 2-h exposure to the pH-matched media control (**Fig. 2B**). Antimicrobials with rapid activity against non-growing or quiescent cell populations often target non-biosynthetic processes, such as lipid membrane homeostasis ^35^. Thus, we hypothesized that the Ms-CFS results in membrane perturbation. As membranes under stress become permeabilized, intracellular ATP and other cytoplasmic contents leak out of the cell into the extracellular milieu. We measured the ratio of extracellular ATP (eATP) to intracellular ATP (iATP) during 1-h exposure of *S. aureus* NRS193 to Ms-CFS. As killing occurred, measured by % survival through viability counts, we observed an increase in eATP/iATP consistent with membrane perturbation and cytoplasmic leakage (**Fig. 2E**).

Several bacteria produce antimicrobial peptides (AMPs) called bacteriocins that antagonize sensitive strains through cell envelope perturbation, including some that form membrane pores ^15^. Subsets of these bacteriocins are also heat stable ^36, 37^. Similarly, boiling of the Ms-CFS with or without the protein denaturing agent DTT did not abolish its antibacterial activity (**Fig. 2F**). Thus, the antibacterial effectors in the Ms-CFS are heat stable. We hypothesized that the effectors could be small heat stable AMPs. However, separation of the Ms-CFS over a 10 kDa filter revealed the fraction <10 kDa (where heat-stable AMPs are expected to elute ^38^) to be completely non-toxic to *S. aureus*, while the fraction >10 kDa remained toxic (**Fig. 2G**). Based on these data, the antimicrobial activity in the Ms-CFS results in *S. aureus* membrane perturbation and rapid killing through a heat-stable effector greater than 10 kDa. While efforts to identify the antimicrobial effector(s) are ongoing, we sought to further characterize the antimicrobial activity in the context of the skin environment.

### The antimicrobial activity of Malassezia secreted products requires an acidic environment

Skin is a highly acidic environment compared to most other body sites, and sebaceous sites in particular can have a surface pH <5 ^39^. This acidity is predicted to contribute to the innate barrier defense of the epidermis against *S. aureus* colonization ^6^. *M. sympodialis* acidifies mDixon throughout 96-h of growth, and all prior experiments with Ms-CFS were adjusted to pH 5.5 with a pH-matched media control (**Fig. 3AB**). As shown in **Fig. 2B**, *S. aureus* viability gradual decreases over 24-h in the media control at pH 5.5. We predicted this was due to the acidity, as *S. aureus* can grow in mDixon at pH 6 over 20-h (**Fig. S3A**). To alleviate this toxicity of the media, we adjusted the Ms-CFS and pH-matched media control to pH 6. Surprisingly, the Ms-CFS toxicity was abolished at pH 6 (**Fig. 3A**). When tested over a pH range from pH 5.4 to pH 6.0, the 2-h Ms-CFS treatment only reduced *S. aureus* viability by greater than 10-fold between pH 5.4-5.6 (**Fig. S3C**). Based on this result, we conclude that the toxicity of the *M. sympodialis* secreted products requires an acidic environment like that which occurs naturally on the skin.

**Figure 3.**
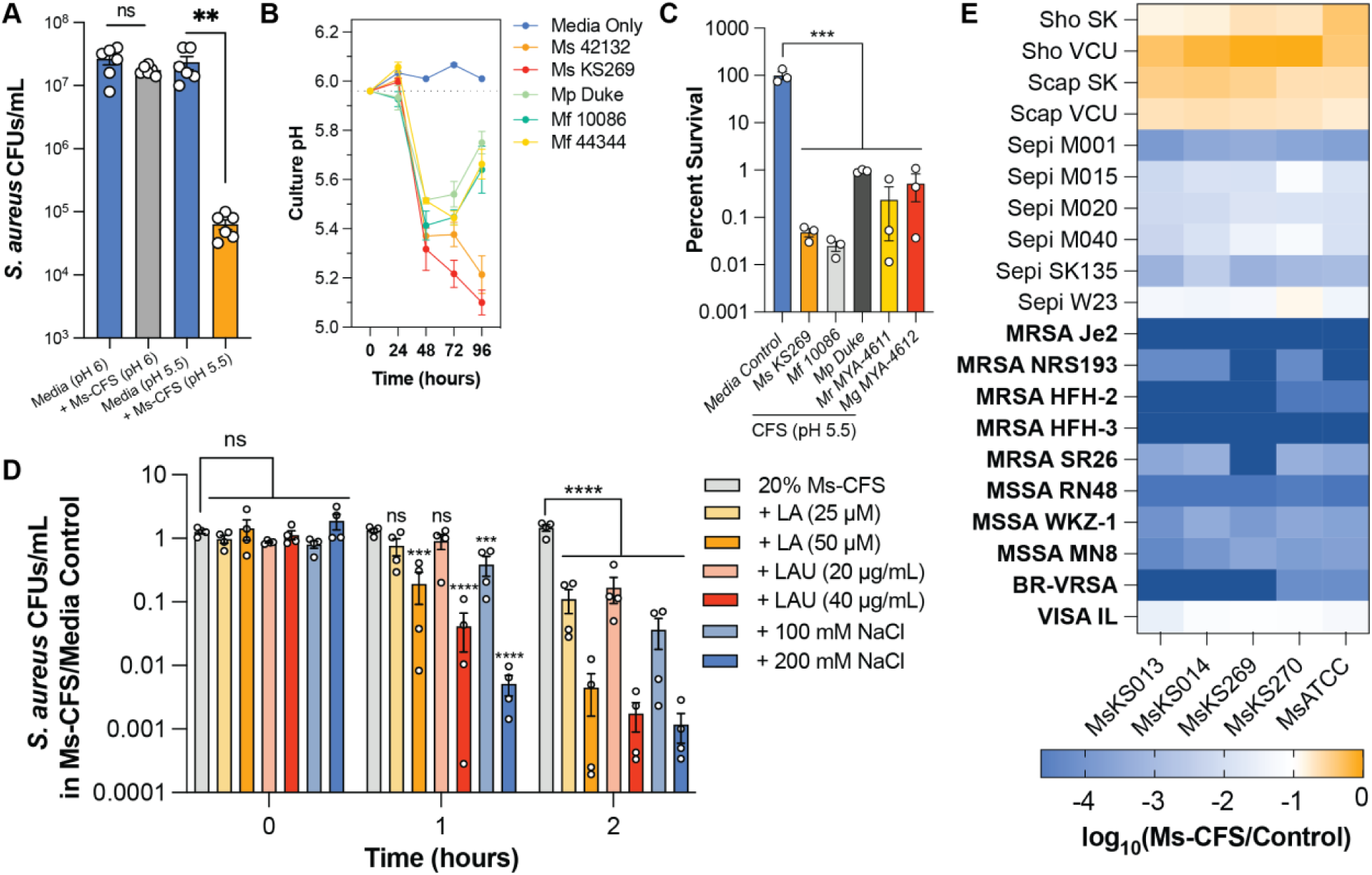
*Malassezia* antimicrobial activity is exacerbated by characteristics of the dermal niche with reduced toxicity to skin-resident staphylococci. **A.** *S. aureus* NRS193 CFUs quantified after exposure to *M. sympodialis* CFS adjusted to pH 6 (grey) or pH 5.5 (orange) or respective pH-matched media controls (blue) after 2-h exposure. Data pooled from two separate experiments, n=6 per group. **B**. The pH of *Malassezia* cultures grown in mDixon over 96-h. Dashed line indicates the initial pH of mDixon prior to inoculation. **C**. Percent survival of *S. aureus* NRS193 in *Malassezia* CFS relative to the media control after 2-h treatment. All CFS at 50% and adjusted to pH 5.5. Data pooled from three separate experiments. Ms: *M. sympodialis*, Mf: *M. furfur*, Mp: *M. pachydermatis*, Mr: *M. restricta*, Mg: *M. globosa*. Strain details in Table S1. One-way ANOVA with Dunnett’s multiple comparisons test, ***: p<0.001**. D**. Log transformed ratio of staphylococcal CFUs from 2-h CFS treatment to Control. Columns correspond to CFS prepared from five *M. sympodialis* strains. Rows present different staphylococcal strains (Sho: *S. hominis*, Scap: *S capitis*, Sepi: *S. epidermidis*, MRSA: methicillin resistant *S. aureus*, MSSA: methicillin sensitive *S. aureus*, VRSA: vancomycin resistant *S. aureus*, VISA: vancomycin intermediate *S. aureus*). A value of 1 indicates a 10-fold reduction in CFUs after 2-h of CFS treatment. The dark navy-blue squares indicates where *S. aureus* CFUs from the CFS-treatment were below the limit of detection (<200 CFUs/mL). **F.** The ratio of *S. aureus* NRS193 CFUs recovered in the CFS condition relative to the media control over 2-h exposure to media or 20% CFS with or without linoleic acid (LA), lauric acid (LAU), or NaCl. Data is from four independent experiments with one-way ANOVA with Dunnett’s multiple comparisons test performed at each time point. ns: not significant, ***: p<0.001, ****: p<0.0001. All error bars indicate SEM.

We hypothesized that previous efforts investigating *Malassezia* interactions with *S. aureus* may not have observed antagonism because of this unique pH sensitive activity ^39–43^. While *M. sympodialis* cultures are naturally acidified by the yeast over 96-h, some *Malassezia spp.* may not acidify the CFS and resulting toxicity may be masked until pH adjustment. When culture pH was monitored over 96-h, two strains of *M. sympodialis* (ATCC42132 and KS269, **Table S1**) both acidify the media from pH 6 to below pH 5.25 (**Fig. 3B**). In contrast, two strains of *M. furfur* (10086 and ATCC44344, **Table S1**) and one strain of *M. pachydermatis* do not acidify the CFS to the same extent by 96-h (**Fig. 3B**). If acidification of the microenvironment is necessary for *Malassezia* antagonism of *S. aureus*, the inability of *M. furfur* and *M. pachydermatis* to inhibit *S. aureus* may be due to differences in pH when cultured on solid media (**Fig. 1**). To test this, we collected and pH adjusted (pH 5.5) CFS from *M. sympodialis*, *M. furfur*, and *M. pachydermatis*. Compared to the pH-matched control, CFS from each species was able to significantly reduce *S. aureus* NRS193 survival after 2-h exposure (**: p<0.01) (**Fig. 3C**). Additionally, CFS collected from the two most prevalent *Malassezia* species on healthy human skin, *M. restricta* and *M. globosa,* showed toxicity to *S. aureus* NRS193 when adjusted to pH 5.5 (**Fig. 3C**). These data suggest that antagonism of *S. aureus* through antimicrobial secreted products is conserved across multiple *Malassezia* species, and that the antimicrobial activity is uniquely active within the acidic microenvironment of the skin.

### Sodium chloride and free fatty acids exacerbate M. sympodialis antimicrobial activity

In addition to its relative acidity, the epidermal surface also contains sodium from sweat at moist sites and free fatty acids (FFAs) from sebum at sebaceous sites that contribute to barrier defense ^5^. We sought to determine if these abiotic features of the skin microenvironment could also impact the toxicity of the yeast secreted products against *S. aureus*. We applied a low dose of Ms-CFS (20% Ms-CFS) that over 2-h does not reduce *S. aureus* viability relative to the pH-matched media control (**Fig. 3D**). We added two subinhibitory concentrations of the antimicrobial FFAs lauric acid (LAU) or linoleic acid (LA) to the 20% Ms-CFS and the control. The addition of either FFA significantly enhanced killing of *S. aureus* NRS193 by the Ms-CFS but not when added to the pH-matched media control (**Fig. 3D**). Differences in FFA abundance between the different media types could contribute to the variation in Ms-CFS potency.

*S. aureus* is halotolerant and thus able to grow at high concentrations of sodium chloride, however exposure to subinhibitory concentrations of sodium chloride has been reported to alter the activity of membrane active compounds, including AMPs ^44–46^. We sought to determine if sodium chloride could impact the toxicity of the Ms-CFS against *S. aureus* NRS193 using 100 mM and 200 mM NaCl in 20% Ms-CFS or the media control. As observed with the addition of FFAs, the addition of sodium chloride significantly increased the toxicity of the Ms-CFS treatment (**Fig. 3D**). Together these data again suggest that the antagonism of *S. aureus* by *M. sympodialis* secreted products is especially potent within the skin microenvironment.

### Skin commensal staphylococci have reduced susceptibility to M. sympodialis secreted products

*S. aureus* is a transient colonizer of most epidermal sites, and typically stable colonization occurs typically only in the anterior nares ^3, 4^. In contrast, several coagulase negative staphylococci are ubiquitous members of the healthy skin microbiota including *Staphylococcus epidermidis, Staphylococcus capitis,* and *Staphylococcus hominis* ^47^. As *Malassezia* are the dominant fungal colonizers of most skin sites, these yeasts likely coexist with commensal staphylococci ^19^. Based on our observation that *S. hominis* can grow adjacent to *M. sympodialis* (**Fig. 1B**), we hypothesized that commensal staphylococci would be able to survive exposure to the yeast secreted products. We treated two *S. homini*s strains, two *S. capitis* strains, six *S. epidermidis* strains, and ten *S. aureus* strains with CFS prepared from five strains of *M. sympodialis* (**Table S1**). Most of the *S. aureus* strains tested showed greater than 100-fold reduction in CFUs after 2-h Ms-CFS treatment compared to the media control (**Fig. 3E**). The only *S. aureus* strain with less than 100-fold reduction in viability was a vancomycin intermediate *S. aureus* (VISA) strain. In contrast to *S. aureus*, *S, hominis* and *S. capitis* were largely unaffected by the Ms-CFS treatment with less than a 10-fold reduction in viability observed. *S. epidermidis* sensitivity was more variable, with some strains experiencing ∼10-fold reduction in viability, while others were phenotypically similar to *S. aureus* (100-fold or greater reduction in viability) (**Fig. 3E**). These data are consistent with the idea that some skin commensal staphylococci have acquired strategies to co-exist with *M. sympodialis*.

### Spontaneous tolerance to growth with M. sympodialis yeast coincides with tolerance to the yeast secreted products

We isolated a surviving strain of *S. aureus* NRS193 (named EVOL-D) that evolved spontaneous tolerance to biofilm growth with *M. sympodialis* from a 6-day mixed colony biofilm (**Fig. S4A**). When EVOL-D and other surviving WT-like isolates from the same mixed biofilm were treated with Ms-CFS or the pH-matched media control, the EVOL-D strain alone could tolerate the treatment with no reduction in viability after 2-h (**Fig. S4B**). This suggests that the toxicity to *S. aureus* is the same whether in co-cultured with *M. sympodialis* yeast cells and during Ms-CFS treatment. These findings also raised the broader question of how pathogens such as *S. aureus* may adapt in response to antagonism by members of the microbiota, including *Malassezia*. To address these questions, we next set out to evolve *S. aureus* tolerance to the Ms-CFS and determine if mechanisms of adaptation are shared between the mixed-biofilm-derived isolate (EVOL-D) and Ms-CFS-derived isolates.

### S. aureus evolves tolerance to M. sympodialis antagonism through mutations in the stringent response regulator Rel

We next applied experimental evolution to select for *S. aureus* tolerance to CFS from *M. sympodialis* grown in mDixon. Here *S. aureus* NRS193 was exposed to the Ms-CFS then surviving cells were recovered in normal culture media (Tryptic soy agar, TSB) before re-exposure to the Ms-CFS again. We repeated this process for 12 passages with three independent populations (Rep A, Rep B, and Rep C). In all three Ms-CFS-exposed populations, we observed increased survival after six passages (P6) (**Fig. 4A**). A control population serially exposed to the pH-matched media control was performed in parallel (Control). After passage 12 (P12), we selected single colonies from each population with phenotypic tolerance to 2-h Ms-CFS exposure (EVOL-A from Rep A, EVOL-B, from Rep B, and EVOL-C, from Rep C) (**Fig. 4B**). The level of tolerance to Ms-CFS in the Ms-CFS-derived strains was similar to that of the mixed-biofilm-derived isolate EVOL-D (**Fig. S4A**). EVOL-A is also more tolerant to mPDB Ms-CFS compared to its NRS193 ancestor and CFS from other *Malassezia* species grown in mDixon (**Fig. S5A, B**), suggesting that the underlying toxicity is similar regardless of media type or *Malassezia* species tested.

**Figure 4.**
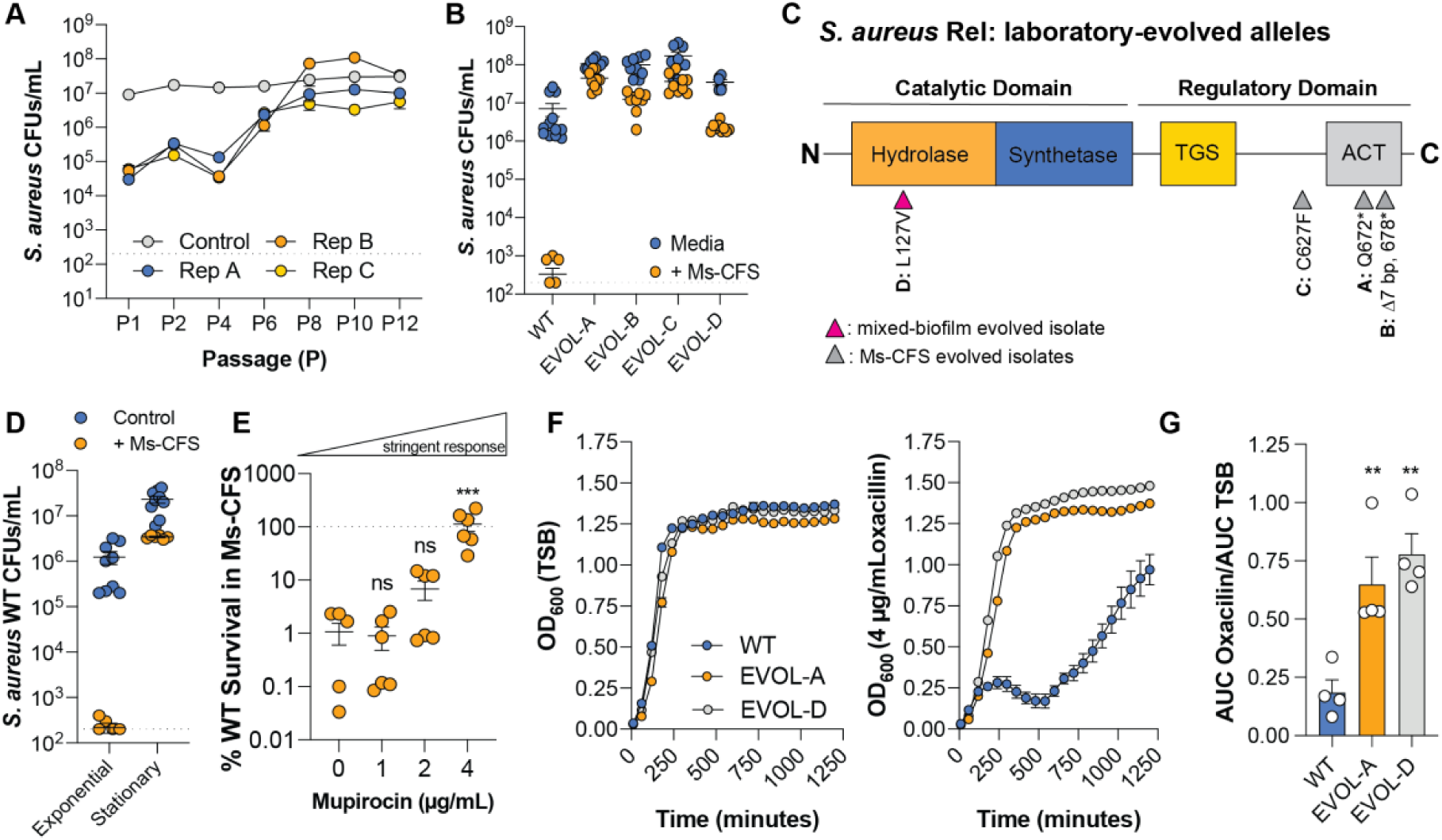
*S. aureus* mutations that activate the stringent response result in tolerance to *Malassezia* antimicrobial activity. **A**. *S. aureus* NRS193 CFUs quantified throughout serially passaging in pH-matched mDixon media (n=1, Control) or *M. sympodialis* CFS (n=3, Rep A-C). **B**. *S. aureus* CFUs from NRS193 wild type (WT) or individual evolved isolates after exposure to media control (Control, blue) or 50% *M. sympodialis* CFS (+CFS, orange). Data pooled from three separate experiments. **C**. Protein domain diagram of *S. aureus* Rel with the approximate locations of the evolved Rel alleles indicated with grey or pink carrots. Asterix indicate a change to a stop codon. **D**. *S. aureus* NRS193 WT CFUs quantified following 2-h 50% *M. sympodialis* CFS treatment when cells are from exponential phase (OD600=∼1) or stationary phase (OD600>4). Data pooled from two separate experiments. **E**. Survival of *S. aureus* NRS193 WT calculated from CFUs after 2-h CFS treatment in the presence or absence of mupirocin. Cells were also pre-exposed to mupirocin for 2-h before treatment. Data pooled from two separate experiments, three independent replicates per experiment. **F**. Growth curves based on OD_600_ for NRS193 WT (blue), EVOL-A (orange), EVOL-D (yellow) in TSB or in TSB with 4 µg/mL oxacillin. Data is representative of four independent experiments. **G.** Area under the curve (AUC) calculation from **F** (n=4). Dashed lines indicate limit of detection, except in E where the dashed line corresponded to 100% survival. **: p<0.01, ***: p<0.001, one-way ANOVA with Dunnett’s multiple comparisons test. TGS: named for its presence in <u>Th</u>rGs, <u>G</u>TPase, and <u>S</u>pot. ACT: named for its presence in bacterial aspartate kinase, chorismite mutase and TyrA.

We next performed whole genome sequencing of the tolerant strains and the ancestor to identify the genetic mechanisms underlying the phenotypic tolerance. Surprisingly, in each of the three Ms-CFS-derived tolerant strains, we identified different mutations in the C-terminal regulatory domain of the stringent response regulator (GTP pyrophosphokinase) Rel (also referred to as Rsh ^48^) (**Fig. 4C**, **Table S2**). In contrast, the mixed-biofilm-derived tolerant isolate, EVOL-D, has a mutation in the N-terminal hydrolase domain of Rel (**Fig. 4C**). Notably, isolates from the final passage (P12) of the Rep A population that were not tolerant to Ms-CFS treatment encoded the ancestral *rel* allele (**Fig. S5C**). In parallel to the experimental evolution of the USA400 WT strain NRS193, we also performed the same experiment with a *S. aureus* USA300 clinical isolate HFH-29568 and again observed evolved tolerance by P12 (**Fig. S5D**). Sequencing from a Ms-CFS tolerant single colony isolated from P12 of this experiment also revealed a mutation in *rel* (**Fig. S5E**, **Table S2**). Together these data suggest that mutations in the stringent response regulator Rel contribute to *S. aureus* survival in the presence of *M. sympodialis* or its secreted products.

### Partial activation of the stringent response results in S. aureus tolerance to M. sympodialis antagonism

The stringent response is typically activated in bacteria in response to starvation, where ribosomes stall due to improper tRNA loading ^49^. Rel interacts with the stalled ribosomes through its TGS domain facilitating a stringent state where the synthetase domain becomes active and consumes GDP or GTP as substrates to produce 5’-diphosphate 3’-diphosphate (ppGpp) or 5’-triiphosphate 3’-diphosphate (pppGpp), respectively. These molecules are referred to as (p)ppGpp or alarmones for simplicity ^50^. In *S. aureus*, *rel* is an essential gene due to the requirement for a functional hydrolase domain that can reduce the pool of (p)ppGpp produced by the synthetase domain of Rel, or by the two small synthetase-only GTP pyrophosphokinases RelP and RelQ ^51, 52^. Accumulation of (p)ppGpp is lethal as it stalls *S. aureus* growth by downregulating protein synthesis, depleting GDP/GTP, and modifying metabolism through the transcriptional repressor CodY ^53^. Mutations that lead to partial activation of the stringent response in *S. aureus* have been described previously to contribute to clinical antibiotic tolerance, however we are not aware of any reports to-date where stringent response activation facilitates tolerance to intermicrobial antagonism ^54^.

The stringent response is active during nutrient deprivation, such as during stationary phase, or when treated with mupirocin during exponential phase ^55–57^. Notably, all prior treatments with Ms-CFS were with *S. aureus* from exponential phase. To determine if stringent response activation promotes *S. aureus* survival to Ms-CFS treatment, we exposed WT NRS193 stationary phase cells and exponential phase cells induced with mupirocin to Ms-CFS. Stationary phase *S. aureus* NRS193 tolerated Ms-CFS treatment with only ∼10-fold reduction in viability compared to the media control (**Fig. 4D**). Similarly, when WT NRS193 cells from exponential phase were treated with increasing levels of mupirocin, the once sensitive cells could now tolerate Ms-CFS treatment (**Fig. 4E**), suggesting activation of the stringent response is sufficient to increase *S. aureus* survival exposed to Ms-CFS.

To test if the stringent response is activated in our evolved tolerant strains, we next measured the ability of EVOL-A and EVOL-D to grow in high levels of the β-lactam oxacillin. When the stringent response is active in MRSA strains, they display homogenous resistance to β-lactams, evident by the ability of the treated population to grow at higher doses of the antibiotic^58^. The minimum inhibitory concentration (MIC) of oxacillin for the NRS193 ancestor was 2-4 µg/mL, while the MIC for EVOL-A and EVOL-D was 16-32 µg/mL oxacillin. Additionally, when cultured in 4 µg/mL oxacillin for 20-h, the Ms-CFS-tolerant strains grow significantly more than the NRS193 ancestor as measured by area under the curve (AUC) (**Fig. 4F, G**). These data support the hypothesis that the stringent response is partially activated under basal conditions in the Ms-CFS-derived and mixed-biofilm-derived tolerant strains. As the mechanism of homogenous resistance during the stringent state relies on increased expression of the penicillin binding protein MecA, we deleted *mecA* in EVOL-A to test if it directly contributes to Ms-CFS tolerance ^59^. While loss of *mecA* reduced growth during exposure to 6 µg/mL oxacillin as expected, the tolerance to Ms-CFS was unaffected (**Fig. S7**).

We quantified pppGpp in a basal state and after stringent state induction with mupirocin^60^. In basal conditions, pppGpp was below the level of detection in all strains, but after induction, EVOL-D showed a significant increase in pppGpp compared to the ancestral WT strain (**Fig. S6AB**). This is consistent with another well characterized *rel* allele (F128Y), also mutated within the hydrolase domain, that leads to stringent response activation and alarmone accumulation ^54^. In contrast, EVOL-A, with a truncated C-terminal domain, did not produce pppGpp at levels different from WT after mupirocin induction (**Fig. S6AB**), most likely because Rel enzymes with C-terminal truncations can no longer interact with the ribosome to sense and respond to amino acid starvation. The observation that all strains had undetectable alarmone levels in a basal state most likely reflects the sensitivity of the assay and the fact that even elevated alarmones in tolerant strains would need to be low enough to not restrict *S. aureus* growth. Based on the phenotyping data, we conclude that the evolved tolerant strains have a constitute, partial activation of the stringent response.

### Laboratory-evolved rel alleles are necessary and sufficient for S. aureus tolerance to M. sympodialis secreted products

The phenotypes of the laboratory evolved *S. aureus* strains with tolerance to *M. sympodialis* CFS are consistent with partial activation of the stringent response. To confirm that the *rel* alleles are both necessary and sufficient for *S. aureus* survival in Ms-CFS, we generated markerless allele swaps. When the ancestral WT *rel* allele was replaced with the EVOL-A *rel* allele (Q672*, WT^Rel-A^) or the EVOL-D allele (L127V, WT^Rel-D^), the strain survives significantly better than the ancestor during exposure to Ms-CFS (**Fig. 5A**). Additionally, introduction of the alleles increases growth in the presence of oxacillin indicating activation of the stringent response and a switch to homogenous β-lactam resistance (**Fig. S8A, B**). Similarly, when the EVOL-A and EVOL-D *rel* alleles are replaced with the ancestral WT *rel* allele (EVOL-A^Rel-WT^, and EVOL-D^Rel-WT^) there is a significant reduction in survival when exposed to Ms-CFS (**Fig. 5A**) and a concomitant reduction in growth in oxacillin (**Fig. S8A, B**). These data indicate that the evolved *rel* alleles are both necessary and sufficient for survival when exposed to *M. sympodialis* antimicrobial products.

**Figure 5.**
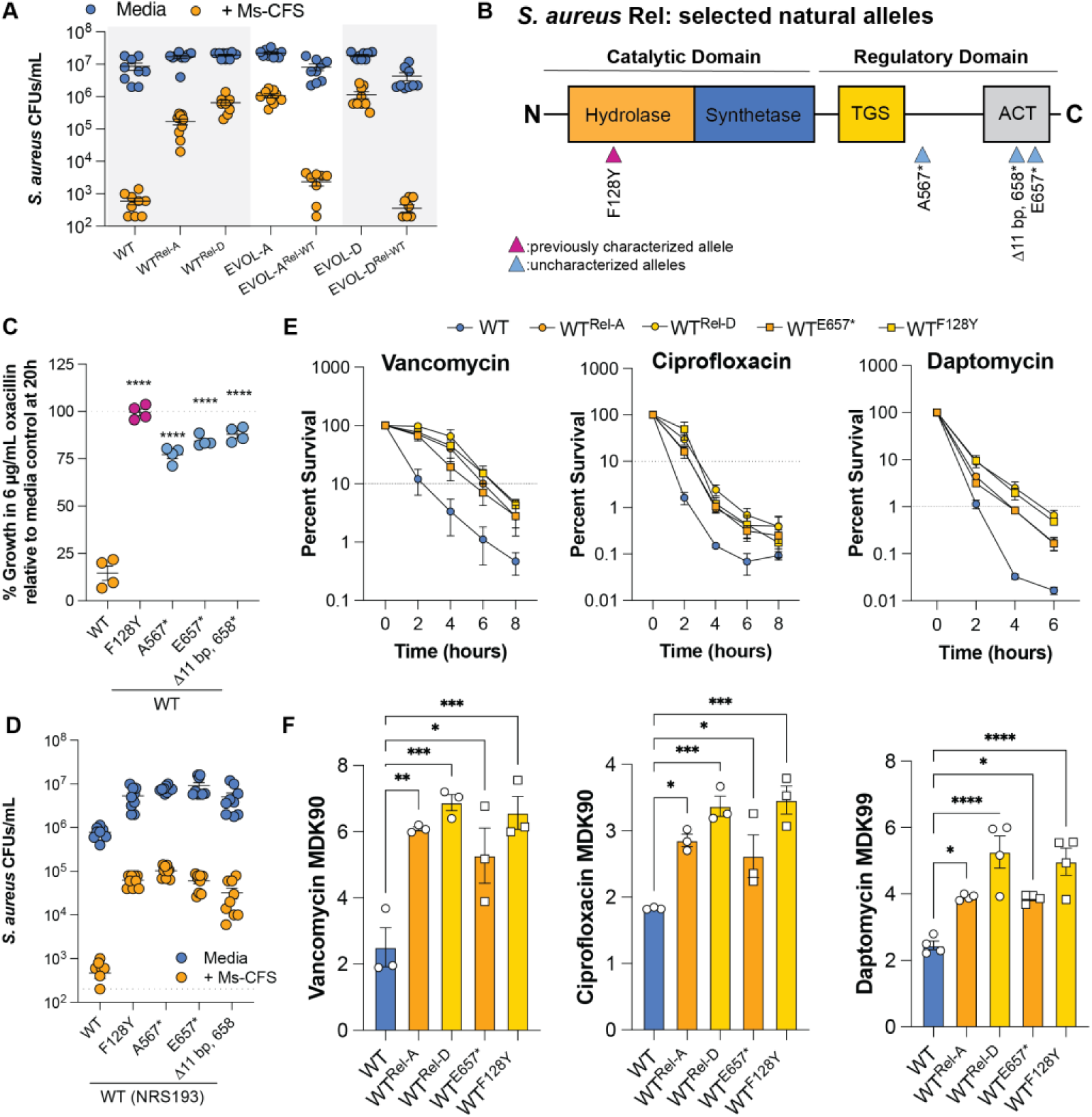
Natural Rel variants that activate the stringent response confer tolerance to *Malassezia* antimicrobial activity and clinical antibiotics. **A**. *S. aureus* CFUs following 2-h CFS treatment (orange) or pH-matched media control (blue) for NRS193 WT and *rel*-allele swap strains. Rel-WT is the NRS193 WT allele, Rel-A is the EVOL-A Q672* allele, and Rel-D is the EVOL-D L127V allele. Data is pooled from three separate experiments, n=9 per strain. **B**. Protein domain diagram of *S. aureus* Rel with the selected natural alleles indicated with pink or blue carrots. **C**. The percent growth of *S. aureus* calculated in 6 µg/mL oxacillin relative to TSB control based on OD_600_ after 20-h. Alleles correspond to the natural Rel alleles in **B**. One-way ANOVA with Dunnett’s multiple comparisons test. ****: p<0.0001. Data pooled from two independent experiments. **D**. *S. aureus* CFUs following 2-h CFS treatment (orange) or exposure to pH-matched media controls (blue). Alleles correspond to the natural Rel alleles in **B**. Data pooled from three independent experiments (n=9 per strain). **E**. Time-kill assays for *S. aureus* strains exposed to vancomycin (1 µg/mL), ciprofloxacin (0.5 µg/mL), or daptomycin (100 µg/mL) over 6 or 8-h. Dashed lines indicate the percent survival at which the minimum duration of killing (MDK) was calculated for each antibiotic. Data represent the mean from three (vancomycin and ciprofloxacin) or four (daptomycin) independent experiments with error bars indicating SEM. **F.** MDK values extrapolated from the linear range of the curves in **E** for each antibiotic. Each point represents the mean of one biological replicate. One-way ANOVA with Dunnett’s multiple comparisons test. *: p<0.05, **: p<0.01, ***: p<0.001, ****: p<0.0001

### Rel alleles from natural S. aureus strains confer tolerance to M. sympodialis antagonism

Mutations in *rel* have been previously identified in clinical strains of *S. aureus* ^54, 61, 62^. One of the first characterized natural Rel mutations is F128Y, which was originally identified in a clinical isolate from a case of persistent bacteremia. Careful study of this allele has shown that it leads to a significant increase in ppGpp consistent with constitutive, partial activation of the stringent response ^54^. Notably, the F128Y mutation is only one position away from the L127V mutation that we identified in EVOL-D (**Fig. 5B**). We therefore sought to determine if natural Rel variants could impact *S. aureus* tolerance to *Malassezia* antimicrobial activity. As expected, based on similarity to EVOL-D, replacing the ancestral alleles with the F128Y allele in NRS193 increases growth in oxacillin (**Fig. 5C**), and increases survival when exposed to Ms-CFS (**Fig. 5D**). This further supports our hypothesis that the Ms-CFS-evolved tolerant strains with mutations in *rel* exist in a partially active stringent state. In the case of F128Y and L127V, we expect these alleles reduce hydrolase activity resulting in a constitutively elevated (p)ppGpp pool ^54^

Natural Rel variants with mutations near or within the C-terminal ACT domain (**Fig. 5B**), have also been described recently ^62^. It is hypothesized that truncation of the ACT domain leads to partial activation of the stringent response by abolishing ligand bind within this region that modulates hydrolase activity ^63–65^. We queried the NCBI Identical Protein Groups database for Rel alleles that specifically had truncations within the C-terminus after the TGS domain. We found several C-terminal truncated Rel alleles fitting these parameters (**Table S3**). From this selection, we chose three natural C-terminal truncation alleles: A567*, E657*, and Δ11 bp that results in a protein truncated at position 658 (Δ11 bp, 658*) (**Figure 5B**). We replaced the WT *rel* allele in NRS193 with these variants and observed all three to increase survival in oxacillin, indicating activation of the stringent response, and increased survival in Ms-CFS by ∼100-fold (**Fig. 5C, D**). The fact that these natural Rel variants lead to *S. aureus* tolerance to *M. sympodialis* antagonism could mean 1) factors that select for activation of the stringent response in *S. aureus* could concomitantly result in *S. aureus* overcoming microbiota-mediated colonization resistance, or that 2) microbiota antagonism can select for *S. aureus* with activation of the stringent response with consequences for antibiotic treatment ^66^.

### Laboratory evolved rel alleles increase S. aureus tolerance to clinical antibiotics

It was previously shown that the F128Y Rel allele results in multidrug tolerance in *S. aureus* ^54^, but other clinical Rel alleles, including a truncation of the C-terminal ACT domain, were subsequently observed to have no impact on antibiotic tolerance ^62^. We sought to determine if the evolved Rel alleles in this study Rel-A (Q672*) and Rel-D (L127V) impacted antibiotic tolerance similar to F128Y. We also included the natural variant E657*, as other natural C-terminal alleles were not previously observed to impact antibiotic tolerance ^62^. Using time-kill assays, we determined the minimum duration of killing (MDK), the gold standard method for quantifying antibiotic tolerance ^67, 68^. As previously reported, the F128Y allele is sufficient to significantly increase tolerance to vancomycin (MDK_90_), ciprofloxacin (MDK_90_), and daptomycin (MDK_99_) (**Fig. 5E, F**) ^54, 68^. Our evolved hydrolase allele, Rel-D, similarly increases tolerance to each antibiotic tested appearing almost indistinguishable from F128Y (**Fig. 5E, F**). The C-terminal alleles also significantly increase tolerance to all three antibiotics, and while the magnitude of tolerance was similar between them, the C-terminal truncations appear to be marginally less tolerant than the N-terminal hydrolase alleles (**Fig. 5E, F**). The multidrug tolerance of the F128Y Rel allele was previously suggested to be the result of extended lag phase, a phenotype that underlies various genetic mechanisms of antibiotic tolerance across diverse bacteria ^67^. Similarly, Rel-A and Rel-D are sufficient to reduce *S. aureus* growth rate compared to the NRS193 ancestor in TSB during the first 4-h of growth (**Fig. S8C**). Whether other physiological aspects of the partially stringent state contribute to antibiotic and Ms-CFS tolerance remains to be determined. These data demonstrate that truncations at the C-terminus of *S. aureus* Rel, as small as 57 amino acids (Q672*), are sufficient for multidrug antibiotic tolerance. Additionally, our findings extend this tolerance from clinical antibiotic to include microbiota-derived antimicrobials.

## Discussion

Recent studies indicate that host resident fungi, despite their relatively low abundance, have major impacts on host health ^69^. For example, the gastrointestinal colonizer *Candida albicans* can induce a protective systemic Th17 response with cross reactivity to other fungal pathogens ^70, 71^. While several studies have identified impacts of the mycobiota on bacterial communities or dysbiosis^72, 73^, there are few studies reporting a specific role for fungi in pathogen colonization resistance ^25, 74^. This is despite examples in other systems, such as the cheese microbiota, where fungi isolated from the polymicrobial community have been observed to impact bacteria growth, survival, and antimicrobial susceptibility ^75, 76^.

Human skin hosts a unique mycobiota that is largely colonized by a single genus of yeast at most epidermal sites of adults ^20^. While *Malassezia* yeasts have been associated with several skin diseases such as pityriasis versicolor and seborrheic dermatitis ^77^, they typically colonize skin without inducing disease. This has led many to hypothesize that *Malassezia* have a mutualistic relationship with the host, where these lipid-dependent yeasts thrive within the lipid-rich niche and may benefit the host by inducing expression of AMPs ^78, 79^. In addition to host-fungal interactions, observations support an important role for *Malassezia* interactions with other microbes that impact host health. For example, in atopic dermatitis, an inflammatory disease often associated with *S. aureus* skin colonization, non-*Malassezia* fungal diversity has been reported to be increased relative to healthy controls ^80^. Additionally, *M. globosa* secretes a protease capable of degrading *S. aureus* biofilms ^25^.

Our results identify a beneficial role for *Malassezia* on human skin, where colonization by *M. sympodialis* significantly reduces the ability of *S. aureus* to colonize the human epidermis (**Fig. 1**). We show that *M. sympodialis,* grown in different culture conditions, secretes antimicrobial product(s) with activity against *S. aureus* that is exacerbated by acidic and fatty acid rich conditions reminiscent of the skin microenvironment (**Fig. 3**). The antimicrobial effector(s) is heat-stable and retained above a 10 kDa molecular weight cut off (**Fig. 2**). Efforts are ongoing to identify the antimicrobial effector(s) and characterize the underlying genes and pathways required for production in *Malassezia*. As molecular approaches to genetically engineer *Malassezia* develop, we aim to test if production of the antimicrobial activity observed *in vitro* is necessary for the observed colonization resistance on the skin, or if additional mechanisms might be important for this interaction. The pH restriction of the antimicrobial activity of the Ms-CFS might explain why *S. aureus* and *Malassezia* spp. are both colonizers of the anterior nares and the skin of atopic dermatitis patients, where the pH is less acidic than most other, healthy epidermal regions ^39, 80–82^.

Despite the identity of the antimicrobial effector(s) remaining unknown, we observed that long term exposure to the *Malassezia* yeasts or their bulk secreted products (Ms-CFS) could rapidly select for stable tolerance to Ms-CFS in multiple *S. aureus* strains (**Fig. 4**). Coincidently, the evolved tolerance to yeast antagonism coincided with tolerance to clinical antibiotics (**Fig. 5**). Antibiotic tolerance is the ability of a microorganism to survive during transient exposure to a high dose of an antibiotic ^68^. Transient, non-heritable antibiotic tolerance in *S. aureus* has been reported to be induced by several host relevant conditions in the absence of clinical antibiotics including human serum ^83^, the macrophage phagolysosome ^84^, and growth as a biofilm ^85^. Additionally, co-culture of *S. aureus* with the pathobiont *C. albicans* in a dual species biofilm was observed to significantly increase *S. aureus* multidrug tolerance compared to *S. aureus* mono-species biofilms ^86, 87^. However, inducers of heritable antibiotic tolerance in *S. aureus* are less well characterized, and it remains an open question if, and to what degree, antagonism by other microbes selects for heritable antibiotic tolerance in the absence of antibiotics. A previous study observed that bacteriocin-producing subpopulations of *S. aureus* could select for vancomycin tolerance in bacteriocin non-producers ^88^. Our *in vitro* studies build upon this literature to support the hypothesis that intermicrobial antagonism has the potential to select for stable, heritable antibiotic tolerance.

*S. aureus* tolerance to *Malassezia* and its antimicrobial products was found to arise from stringent response activation caused by mutations in the master regulator Rel. The stringent response is a well-documented mechanism of multidrug tolerance in *E. coli*, where its activation during biofilm growth contributes to its inherent antibiotic tolerance ^89^. Where in Gram-negative bacteria the stringent response is regulated by two separate proteins, RelA and SpoT, Gram-positive bacteria, such as *S. aureus*, encode a single RelA/SpoT homolog (RSH) superfamily protein Rel ^52^. There have been multiple reports of *S. aureus* and *Enterococcus faecium* clinical isolates with mutations in Rel ^90, 91^, of which mutations within the N-terminal hydrolase domain have been empirically linked to antibiotic tolerance for both species by partially activating the stringent response ^54, 91^. Mutations within the C-terminus have also been identified in distinct clinical isolates of *S. aureus*, yet it remains unknown if these mutations also contribute to multidrug tolerance ^62, 90^. Querying publicly available assembled *S. aureus* genomes, we identified numerous Rel alleles with truncations in the C-terminal regulatory domain (**Table S3**), further highlighting that these alleles occur naturally within *S. aureus* populations. When introduced in the USA400 strain NRS193, all three C-terminal truncation alleles tested conferred tolerance to *M. sympodialis* secreted products (**Fig. 5**). One of these selected for further analysis (E657*) also resulted in multidrug tolerance when introduced into *S. aureus* NRS193. Thus, mutations across the Rel protein sequence allow for tuning of the stringent response to facilitate antimicrobial tolerance, and microbe-microbe interactions may play an important role in selecting for a semi-stringent state. Future work will investigate the downstream mechanisms that allow partial activation of the stringent response, or a semi-stringent state, to confer such broad tolerance to antimicrobials.

Microbiota-mediated colonization resistance against pathogens is well-documented, yet how pathogens may adapt in response to this antagonism has remained largely unexplored. In the case of opportunistic pathogens that occur within the commensal microbiota, pre-adaptation to microbiota competition can select for traits that are beneficial during infection, such as through the production of capsule in *Streptococcus pneumoniae* ^92, 93^. For pathogens invading a new niche, adaptation to antagonism by the microbiota is likely dependent on the rarity of the invader, microbiota density, micro-niche structure, and dynamic environmental selective pressures ^93, 94^. This complexity makes is difficult to empirically assess how the microbiota shapes pathogen adaptation, but natural examples suggest this is the case. First, the epidemic USA300 lineage of *S. aureus* acquired the arginine catabolic mobile element from the commensal *S. epidermidis* that contributes to its survival on skin ^95^. Second, the evolution of methicillin resistance in *S. aureus* is proposed to have been selected for in hedgehogs colonized with β-lactam producing dermatophytes prior to the use of these antibiotics in the clinic ^96^. While additional studies are necessary to confirm that *Malassezia* shapes *S. aureus* adaptation to the epidermal niche, our results suggest that natural variation, evident in *S. aureus* clinical strains and genomic data, is sufficient for tolerance to a broad range of antimicrobials. This poses the question of how broad tolerance mechanisms could evolve as the result of intermicrobial antagonism with consequences for future antibiotic treatment. Additionally, how *Malassezia* shapes the composition of the skin microbiota remains an open question, but the observation that *S. hominis* and *S. capitis* can tolerate growth in *M. sympodialis* secreted products suggest that mechanisms of tolerance may have evolved that allow these organisms to co-colonize human skin (**Fig. 1**, **Fig. 3**). Collectively our findings identify potent novel antimicrobial activity of the skin mycobiota against *S. aureus*, as well as reveal mechanisms by which pathogen adaptation to the mycobiota confers tolerance to clinical antibiotics.

## Acknowledgments

We thank Dr. Joe Heitman for providing the *Malassezia* strains and Dr. Karen Guillemin for providing the *E. coli* Nissle strain. We also thank Dr. Kristin Kohler for critical feedback on the manuscript. This work was carried out with the support of the University of Oregon GC3F core facilities. Funding: CHK (L’Oréal USA For Women in Science Fellowship, Helen Hay Whitney Foundation Fellowship), TJS (National Institutes of Health grants 1P01GM125576, F32DK124033, K99DK137017), MFB (National Institutes of Health grants R35GM133652, R21AI173839), RMC (Sir Henry Dale Fellowship jointly funded by the Wellcome Trust and the Royal Society, 104110/Z/14/A; a Lister Institute Research Prize Fellowship 2018).

## Author Contributions

Conceptualization: C.H.K and M.F.B.; Methodology: C.H.K. and M.F.B.; Investigation: C.H.K., S.L., T.J.S, and R.M.C.; Writing – Original Draft: C.H.K. and M.F.B.; Writing – Reviewing & Editing: S.L., T.J.S., and R.M.C.; Funding Acquisition: C.H.K. and M.F.B.

## Supplemental Information

Document S1: Figures S1-S8, Table S1-S4

## Materials and Methods

### Bacterial and yeast culture conditions

*S. aureus* and other staphylococci were maintained on tryptic soy agar (TSA, Difco) or broth (TSB, Difco) at 37°C. Unless otherwise noted, staphylococci were cultured overnight to stationary phase (∼16-h) and then subcultured in TSB 1:10 for 1-h to an OD_600_ of ∼1 corresponding to exponential phase. The wild type (WT) *S. aureus* strain used in this study is the ST1, USA400 isolate NRS193. This isolate was selected because it did not originate from a skin infection, is available through BEI Resources, and is closely related to the USA400 reference strain MW2 (**Table S1**). The sequenced and annotated genome for NRS193 has been deposited in NCBI as part of this study (Accession Number: TBD). *E. coli* strains were maintained in LB (Sigma) with antibiotic as needed: 100 µg/mL ampicillin or 25 µg/mL chloramphenicol. *S. aureus* was cultured with 10 µg/mL chloramphenicol when necessary.

*Malassezia* species were maintained on mDixon media (36 g/L Malt Extract (Sigma), 20 g/L Ox-bile (Sigma), 10 mL/L Tween40 (Sigma), 6g/L peptone (from casein and other animal proteins, Sigma), 2 mL/L glycerol (Sigma), 2 mL/L oleic acid (technical grade, Sigma), 15 g/L agar (Bacto), adjusted to pH 6 with HCl). Due to supply chain issues, some experiments were performed with Ox-Bile sourced from Hi-Media, and this was used at 10 g/L. For *M. globosa* and *M. restricta*, Tween60 (Sigma) replaced Tween40 and 0.5 g/L glycerol monostearate (Spectrum Chemical) was added. All *Malassezia* were grown at 30°C. Unless otherwise noted the strain of *M. sympodialis* used is KS269 ^97^ (**Table S1**). When noted mPDA (modified potato dextrose agar) was utilized and contains PDA (Sigma) with 4g/L Ox-bile (Sigma, or 2 g/L Hi-Media), 4 mL/L Tween60 (Sigma), and 1 mL/L Tween20 (Fischer) as previously described ^30^. For modified potato dextrose broth (mPDB), the recipe was the same as mPDA except mPDB (Sigma) base was used. The synthetic sebum media is based on SSM9PR media described previously with 0.1% synthetic sebum (Scientific Services S/D Inc.) ^98^. It was prepared as follows 11.28g/L M9 Salts (Sigma), 10 g/L dextrose, 10 g/L casamino acids (Research Products International), 2 mM MgSO_4_, 0.1 mM CaCl_2_, 1 mg/L calcium pantothenate (Acros), 1 mg/L thiamine-HCL (Cayman Chemical), 1 mg/mL nicotinamide (Sigma), 1 mL/L synthetic sebum (Scientific Services S/D Inc.), 3 mL/L Tween80 (Sigma). Note that Tween80 and synthetic sebum were first mixed 3:1 and autoclaved separately. Ms-CFS collected from the synthetic sebum was after 120-h of *M. sympodialis* KS269 growth.

### Adjacent colony assay and mixed colony biofilms

For adjacent colony assays, *Malassezia* species were collected from a 96-h culture from 30°C and washed in fresh mDixon media. The OD_600_ was adjusted to 1 and 10 µL was spotted in triplicate on mDixon agar and incubated at 30°C for 72-h. *S. aureus* or *S. hominis* SK119 were cultured overnight in TSB and subcultured as described above before cells were pelleted and resuspended in mDixon. The OD_600_ was adjusted to 0.1 in mDixon, and 10 µL was spotted adjacent to the 72-h *Malassezia* colonies. Plates were returned to 30°C for 24-h and then photographed. The protocol was similar for *E. coli*, except it was grown in LB.

For mixed biofilm assays *S. aureus* and *Malassezia* were prepared similarly as above but were adjusted to OD_600_ of 0.2 and 2 in mDixon respectively. For the co-culture condition, *S. aureus* and *M. sympodialis* were mixed 1:1 and 15 µL was spotted on mDixon agar. For monoculture conditions, *S. aureus* was mixed 1:1 with mDixon (final OD_600_=0.1) and 15 µL was spotted on mDixon agar. Plates were incubated for 6 days at 30°C, after which entire colony biofilms were scraped up and resuspended in 500 µL of mDixon. After vigorous vortexing (maximum power, 1 min.) to disrupt aggregates, the suspension was serially diluted to plate for *S. aureus* viable counts on TSA at 37°C. When performed with mPDA, the protocol was the same except mDixon broth and agar were replaced by mPDB and mPDA respectively.

### Co-colonization of human skin biopsies

Healthy human skin biopsies were sourced as NativeSkin Access 11mm biopsies from Genoskin (Salem, MA, USA). Genoskin represents the widest network of IRB-approved skin sourcing across Europe and the United States. The biopsies are prepared from discarded tissue following surgery and collected with patient informed consent in respect of the Declaration of Helsinki and approval of the French Ministry of Research and Higher Education and French Ethics Committee. The use of NativeSkin human skin biopsies falls within the category of ‘unidentifiable biospecimens obtained from a provider’ and are not considered human subject research. For this study biopsies are embedded in a matrix and maintained with culture media that is free of antibiotics and antifungals. Biopsies remain viable at 37°C with 5% CO2 for up to 10 days according to the provided protocols.

*M. sympodialis* KS269 cells were grown at 30°C for 72-h in mDixon, washed twice with sterile PBS and then resuspended in PBS with 10 µg/mL calcofluor white (CFW) (fluorescent brightener 28, MP Biomedicals) for 30 min. at room temperature with agitation. We confirmed this concentration of CFW was nontoxic as yeast viable counts do not decrease after labeling (**Figure S2A**). Yeast were pelleted, resuspended in PBS, and enumerated with a hemocytometer. Based on hemocytometer counts, 5×10^5^ yeast were inoculated per biopsy in 10 µL of PBS. Control biopsies received 10 µL of PBS. After 6-days of incubation at 37°C, 5% CO_2_, *S. aureus* expressing GFP was inoculated as follows. *S. aureus* constitutively expressing GFP on a plasmid was cultured in TSB with 10 µg/mL chloramphenicol to stationary phase and then subcultured 1:10 for 1-h in the same conditions. Cells were washed twice in PBS and then resuspended in PBS with 10 µg/mL chloramphenicol at an OD_600_=0.5 that corresponds to ∼5×10^6^ CFUs/10 µL inoculum. Each biopsy was inoculated with 10 µL *S. aureus* suspension. After 24-h at 37°C, 5% CO_2_, one biopsy from each group was removed from the transwell and O-ring with sterile tweezers and inverted into a Mat Tek glass bottom imaging dish (35 mm dish, No. 1.5 uncoated, 10 mm glass diameter) with 200 µL of PBS. Biopsies could then be imaged on an inverted Nikon CSU-W1 SoRa spinning disk microscope using Nikon Elements software. The skin surface is auto-fluorescent with 561 nm excitation which was used to locate the skin surface. The CFW-labeled yeast were visible with excitation at 405 nm and *S. aureus*^GFP^ was visible with excitation at 488nm. Imaging was performed with a 40X water immersion objective. Max projection images were generated in Nikon Elements.

To collect CFUs from the skin biopsies, the skin was removed from the transwell and O-ring with sterile tweezers. Sterile scissors were used to remove most of the hypodermis and cut the skin into four pieces. Skin pieces were placed in 1 mL sterile PBS and vortexed at maximum speed for 3 min. to disrupt the attached microorganisms. Biopsies were then shaken at 250 rpm for 2-h after which they were again vortexed at maximum speed for 2 min. The disrupted skin is allowed to settle briefly, and the PBS is removed, diluted, and plated on TSA for *S. aureus* CFUs. The experiment was repeated with biopsies from three separate donors.

### Cell-free supernatant preparation and treatment

Five to 10 large *M. sympodialis* colonies from a 72-h mDixon agar plate were used to inoculate mDixon broth (typically 12.5 mL in broth in 125 mL flask). Cultures were incubated at 30°C with shaking at 200 rpm for 96-h. Yeast were pelleted for 2 min at 5000 rcf and the supernatant decanted. The pH where noted was adjusted to the desired pH with HCl or NaOH. A pH matched media control was also generated. Both media control and supernatant were filter-sterilized with a syringe tip filter through a 0.22 µm MCE filter (Milipore). This sterilized cell -free supernatant (CFS) and control media were stored at 4°C and before use in experiments were warmed in a 37°C water bath for 10 min.

Unless otherwise noted, all experiments were performed with 50% CFS that was generated by mixing the CFS 1:1 with the pH matched media control. *S. aureus* and other staphylococci were cultured as described above in TSB before cells were pelleted and resuspended in mDixon (pH 6). *S. aureus* and other staphylococci were inoculated into the 50% CFS or pH-matched controls at a final OD_600_=0.02. After treatment, cells were serially diluted and plated on TSA to calculate CFUs/mL.

### Treatment of S. aureus-colonized human skin biopsies

Healthy human skin biopsies were sourced as NativeSkin Access 11 mm biopsies from Genoskin and maintained as described above according to the manufacturer’s instructions. *S. aureus* expressing red fluorescent protein (RFP) on a plasmid (*S. aureus^RFP^)* and shown to be sensitive to *M. sympodialis* CFS *in vitro* (**Fig. S3B**), was cultured overnight in TSB with 10 µg/mL trimethoprim and subcultured to as described above. Cells were washed and resuspended in sterile PBS. In 12 µL, ∼2.5×10^7^ cells were inoculated onto the surface of the skin biopsies and incubated at 37°C, 5% CO2 for 24-h. After 24-h, control biopsies were treated with pH-matched mDixon (+ vehicle) and the test biopsies (+ exoproducts) were treated with 50% CFS collected from *M. sympodialis in vitro* cultures as described above. After 24-h of treatment, the biopsies were removed from the transwells and disrupted as described above to enumerate the recovered *S. aureus* CFUs/biopsy as a metric of colonization. The experiment was repeated with biopsies from three separate donors.

### ATP measurements

Intracellular and extracellular ATP, iATP and eATP respectively, were quantified using the BacTiter Glo^TM^ Microbial Cell Viability Assay (Promega). *S. aureus* was cultured in TSB as described above and adjusted to a final OD_600_=0.1 in 5 mL of *M. sympodialis* 50% CFS or pH matched mDixon control and incubated at 37°C, 250 rpm. At 0, 15, 30, and 60 min., 1 mL aliquots were centrifuged for 1 min. at 5,000 xg and 0.01 mL aliquots were used for serial dilutions of viable counts. The supernatant was transferred to a new tube for quantification of eATP, and cells were resuspended in mDixon for quantification of iATP. The BacTiter Glo^TM^ reagent was prepared according to manufacturer’s protocol and mixed 1:1 with the supernatant or cell suspension. Luminescence was measured using a BioTek Synergy H1 monochromator-based multi-mode microplate reader (BioTek) with black-walled, glass-bottom 96-well plates (Corning) with 0.8s integration time per well. Luminescence for iATP was normalized to OD_600_.

### Processing of cell-free supernatant

*M. sympodialis* CFS was collected from 96-h monocultures as described above. Five mLs of undiluted CFS or mDixon media were separated across a 10 kDa membrane (Thermo, Pierce) by centrifuging in a fixed-angle rotor for 5 min. at 7,000 xg based on the manufacturer’s protocol. The <10 kDa and >10 kDa fractions were brought to 5 mL volume with sterile water. The pH was adjusted to pH 5.5 with HCl or NaOH and filter sterilized as described above. For boiling experiments, 1 mL of *M. sympodialis* 50% CFS pH 5.5 or mDixon pH 5.5 were incubated at 95C for 15 min. in the presence or absence of 1 mM DTT (dithiothreitol, Thermo). *S. aureus* was exposed to the CFS or media control for 2-h as described above, and viable counts enumerated.

### Serial passaging experimental evolution

*S. aureus* (NRS193 (WT) or HFH-29568) were cultured in TSB as described above before the OD_600_ was adjusted to 0.02 in 50% *M. sympodialis* CFS (3 populations) or pH-matched mDixon (1 population). After 8-h of exposure, a 10 µL aliquot was removed to enumerate viable counts from the four populations and the remaining volume of 190 µL was inoculated into 5 mL of TSB to facilitate growth of surviving cells overnight at 37°C. The following day a portion of the recovered population was cryopreserved, the subculture in TSB to exponential phase performed as described above, and then inoculated in 50% CFS (the three separate populations) or pH-matched mDixon at OD600=0.02 for 8-h. This was repeated for 12 passages. From the viable counts surviving passage 12, individual colonies were isolated and tested for their tolerance to 2-h exposure to the CFS. One representative tolerant isolate was selected from each of the CFS-exposed populations of the USA400 WT strain NRS193 (EVOL-A, EVOL-B, and EVOL-C).

### S. aureus genomic DNA preparation

*S. aureus* genomic DNA was prepared using the Qiagen Blood and Tissue kit with the following modifications. From a freshly inoculated TSA plate, 10-20 *S. aureus* colonies were inoculated into 0.1 mL TE buffer with 20 µg/mL lysostaphin (Sigma) and incubated at 37°C for 15-30 min. or until visibly cleared. The lysis buffer for gram positive bacteria described in the Qiagen Blood and Tissue protocol was prepared 2X and without lysozyme (40 mM Tris-HCl pH 8, 4 mM Na2EDTA, 2.4% (v/v) triton X-100) and diluted to 1X by mixing with the lysostaphin-treated cells. The Qiagen protocol was then followed as described.

### Whole genome sequencing and variant calling

Whole genome sequencing, assembly, and annotation of *S. aureus* NRS193 (Accession: TBD) was performed by Plasmidsaurus. Genomic DNA was prepared as described above. Long-read sequencing was performed using Oxford Nanopore Technologies with V14 library preparation and R10.4.1 flow cells. Genome Assembly was performed using Filtlong v0.2.1, Miniasm v0.3, Flye v2.9.1, and Medaka v1.8.0. Genome annotation was performed with Bakta v1.6.1.

Genome sequencing of the evolved strains was performed using SeqCenter Illumina Whole Genome Sequencing. Genomic DNA was prepared as described above and the Illumina DNA Prep kit with bead-based tagmentation was utilized for library preparation. The Illumina sequence data (fastq) was used with the NRS193 reference genome to identify variants using *breseq* with default settings ^99^

### Mupirocin treatment

*S. aureus* was cultured overnight in TSB and then subcultured 1:20 in TSB with 0, 1, 2, or 4 µg/mL mupirocin (Selleck Chemicals) for 2-h. Cells were centrifuged, washed with mDixon, and inoculated into *M. sympodialis* 50% CFS or pH-matched media control with 0, 1, 2, or 4 µg/mL mupirocin at OD_600_=0.02. After 2-h treatment, *S. aureus* cells were serially diluted to enumerate viable counts.

### Bacterial growth curves

Growth curves were performed using a BioTek Synergy H1 monochromator-based multi-mode microplate reader (BioTek) with non-treated, sterile, polystyrene 96-well plates with lids (Celltreat). Unless otherwise noted, growth curves were performed for 20h with OD_600_ measured at 10 min. intervals with incubation at 37°C and continuous orbital shaking between intervals. Each experiment included three to four technical replicates for each condition. For all conditions, *S. aureus* was inoculated at an OD_600_=0.02. The control condition for all experiments was tryptic soy broth (TSB)(Difco). Oxacillin (Sigma) was added as 4 or 6 µg/mL and diluted from a working stock of 50 mg/mL in water. The final volume per well was 150 µL. Data was collected using the BioTek Gen5 software and analyzed in Microsoft Excel and GraphPad Prism 10. Area under the curve (AUC) was calculated in GraphPad Prism 10 with the baseline at Y=0. Peaks were ignored if the peak height was less than 10% of the distance from the Y minimum to Y maximum and all peaks must be above the baseline.

### Genetic manipulation of S. aureus

For allelic replacement and in-frame deletions we generated constructs using the pIMAY system as previously described ^100^. All plasmids and primers used are in **Table S4**. Briefly, the *mecA* knockout construct was generated by amplifying 500-1000bp upstream and downstream of *mecA*. Primers P1 and P2 amplified the 5’ flank and primers P3 and P4 amplified the 3’ flank, where P3 includes homology to P2 (Phusion polymerase, NEB). Overlap extension PCR with primers P5 and P6 set ∼100 bp inside of P1 and P4, respectively, was performed to stitch together the flanks. For the *rel* allele swaps, the allele of interest was amplified from genomic DNA using Rel_swap_P1 and Rel_swap_P2 primers. PCR fragments and the fused construct were column purified (Zymo DNA Clean-Up and Concentration kit). The purified construct was phosphorylated with T4 polynucleotide kinase (NEB) according to the manufacturer’s protocol. pIMAY, purchased from Addgene, was prepared from *E. coli* DH5a using the ZymoPure II Plasmid Midiprep kit, digested with SmaI (NEB), and treated with shrimp alkaline phosphatase (rSAP)(NEB) according to the manufacturer’s protocol. The purified fusion construct was ligated into SmaI-digested pIMAY with Quick Ligase (NEB) and transformed into chemically competent *E. coli* DC10B ^100^. Colonies were isolated on 25 µg/mL chloramphenicol and the insertion into pIMAY confirmed with primers IM51 and IM51. Confirmed plasmids were midi-prepped from a 100 mL culture as described above and further concentrated using Novagen Pellet Paint Co-precipitant (Millipore Sigma).

Electrocompetent *S. aureus* strains were generated as previously described ^101^ and transformed with ∼5 µg of the pIMAY construct using the BioRad Gene Pulser Xcell electroporation system and 1 mm disposable electroporation cuvettes (VWR Signature). The pulse was 21kV/cm, 100 Ω, and 25 mF and cells were immediately resuspended in 1 mL of TSB + 500 mM sucrose and incubated at 28°C or 30°C for 1-2-h before plating on TSA with 10 µg/mL chloramphenicol and incubating at 28°C for 48-h. Large colonies were resuspended in 500 µL of TSB and 10 µL streaked on a new TSA plate with 10 µg/mL chloramphenicol and incubated at 37°C overnight to initiate integration onto the chromosome. Large colonies were screened with colony PCR (Phusion polymerase, NEB) for integration through the 5’ flank using primers P1 and P6. Positive colonies were grown overnight at 28°C in TSB without antibiotic and then subcultured 1:10 in TSB with 1 µg/mL anhydrotetracycline hydrochloride (aTC) (Sigma) at 18C for ∼6-h before 10 µL was quadrant streaked on a TSA plate with 1 µg/mL aTC and incubated at 28°C for 48-h. Single colonies were patched on TSA with 1 µg/mL aTC or TSA with 10 µg/mL chloramphenicol. Only colonies that grew on aTC and not chloramphenicol were screened for the gene deletion with primers P1 and P4 and internal primers.

For the introduction of the natural *rel* variants, the construct generated above with the WT *rel* allele in pIMAY (pRel-swap, Table S2) was used for site directed mutagenesis with the Q5-Site-Directed-Mutagenesis kit (NEB) according to the manufacturer’s instruction. The mutagenesis primers were designed using the NEBaseChanger version 1.3.3. Mutagenesis was confirmed with either whole plasmid sequencing or Sanger sequencing. Mutated plasmids were transformed into *E. coli* DC10B and *S. aureus* as described above. Allele replacement was confirmed with Sanger sequencing in *S. aureus*.

### Identification of natural Rel alleles

To identify natural Rel alleles that resemble those in the CFS-derived tolerant strains, we queried the NCBI Identical Proteins Database. We used ‘GTP pyrophosphokinase’ as the search term and *Staphylococcus aureus* as the organism. To eliminate sequences corresponding to RelP and RelQ, we limited out results to a sequence length between 300 and 1000 amino acids. This resulted in 283 unique protein sequences. Based on the methionine annotated as the start codon, the majority of Rel sequences are 729 or 736 amino acids in length. All 283 FASTA files were downloaded and aligned in SnapGene. Two sequences were excluded that appeared to be misannotated as *S. aureus.* We focused on alleles with C-terminal truncations and compiled those in **Table S3**. These truncated alleles have one or two genomes corresponding to each sequence. Following this query, a study was published where C-terminal mutations like those observed in the CFS-derived strains were observed in clinical isolates ^62^.

### Minimum inhibitory concentrations and time-kill assays

Minimum inhibitory concentration for oxacillin were determined following protocols previously described ^54^. Briefly, *S. aureus* grown overnight in TSB and subcultured to exponential phase were inoculated into 0-128 µg/mL oxacillin at 2-fold dilutions to an initial OD_600_=0.02 in a 96-well plate. The final volume per well was 150 µL. After 24-h at 37°C, the minimum inhibitory concentration is reported as range from the highest concentration where *S. aureus* grows to the lowest concentration where there is no growth.

Time-kill assays were performed as previously described ^54^. Briefly, *S. aureus* was grown overnight in Mueller-Hinton Broth (MHB) (Sigma) and subcultured 1:10 for 1-h at 37°C in the same media. *S. aureus* was inoculated into MHB with the following antibiotics: 1 µg/mL vancomycin (Cayman Chemical), 100 µg/mL daptomycin (Selleck Chemical), or 0.5 µg/mL ciprofloxacin (Sigma). These antibiotics were selected because they had previously been shown to have reduced activity against *S. aureus* expressing the F128Y Rel allele in contrast to other antibiotics that had no differences in sensitivity ^54^. Aliquots were taken at 0, 1, 2, 4, 6, and 8-h as necessary to enumerate *S. aureus* viable counts by serial dilution and plating on TSA. The minimum duration of killing was calculated for each independent experiment using the data points within the linear range and applying the linear regression and interpolation functions in GraphPad Prism as previously described ^54^.

### Quantification of pppGpp

*S. aureus* strains were grown overnight in low-phosphate chemically defined media (CDM) ^102^ at 37°C. Cultures were diluted to an OD_600_ of 0.1 and grown for 2-h prior to the addition of 3.7 MBq of [^32^P]H_3_PO_4_ and incubation for a further 3-h at 37°C. Cultures were subsequently normalized for absorbance and to one set the stringent response was induced by the addition of 60 µg/mL mupirocin. Cells were incubated at 37°C for 30 min, before being recovered by centrifugation (17,000 × g for 5 min) and suspended in 100 μL of 2 M formic acid. Cells were subjected to three freeze/thaw cycles and debris removed by centrifugation (17,000 × g for 5 min) before the lysate was filtered through a 3 kDa spin column. Ten microliters were subsequently spotted on PEI-cellulose F thin-layer chromatography (TLC) plates (Merck Millipore), nucleotides separated, and TLC plates developed using a 1.5 M KH_2_PO_4_ pH 3.6 buffer. The radioactive spots were visualized using an FLA 7000 Typhoon PhosphorImager, and data were quantified using ImageQuantTL software.

### Statistical Analysis

Statistical analyses were performed using Prism 10 (GraphPad). Figure legends include details on statistical test, sample number, donor number, and p-value. For skin biopsy experiments, each data point (n) represents a single biopsy with data points color-coded by donor. For bacterial experiments (n) represents biologically independent samples with independent experiments noted. P values: p >/= 0.5 not significant (ns), p<0.05 (*), p<0.01 (**), p<0.001 (***), p<0.0001 (****).

