## Supplemental Data: FigS1-8, TableS1-4 for "Adaptation to skin mycobiota promotes antibiotic tolerance in *Staphylococcus aureus*"

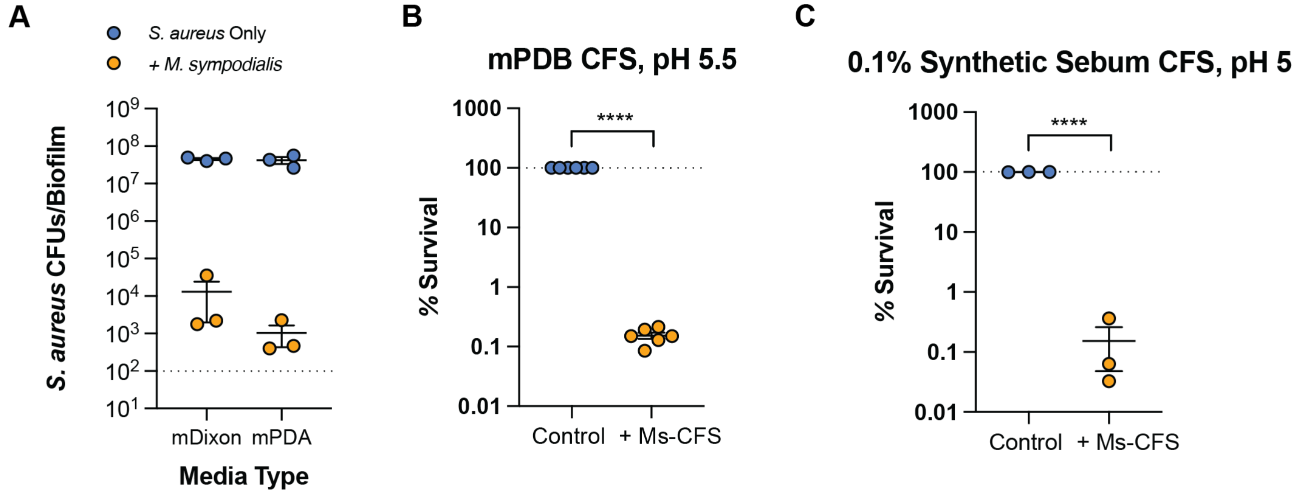


**Figure S1.** ***M. sympodialis in vitro*** **antimicrobial activity with various media. A.** *S. aureus* NRS193 CFUs recovered from 6-day mixed colony biofilm on mDixon or mPDA agar. Data from three experiments. Dashed line is the limit of detection 100 CFUs/mL. *S. aureus* survival after 24-h exposure to **B)** 50% CFS from *M. sympodialis* grown in mPDB (pH 5.5) (n=6) or **C)** 100% CFS from *M. sympodialis* grown in 0.1% synthetic sebum media (pH 5)(n=3) relative to the media control (set to 100% survival). Unpaired t-test, ****: p<0.0001. Whiskers are Min to Max with all points shown. LOD: limit of detection (200 CFUs/mL).

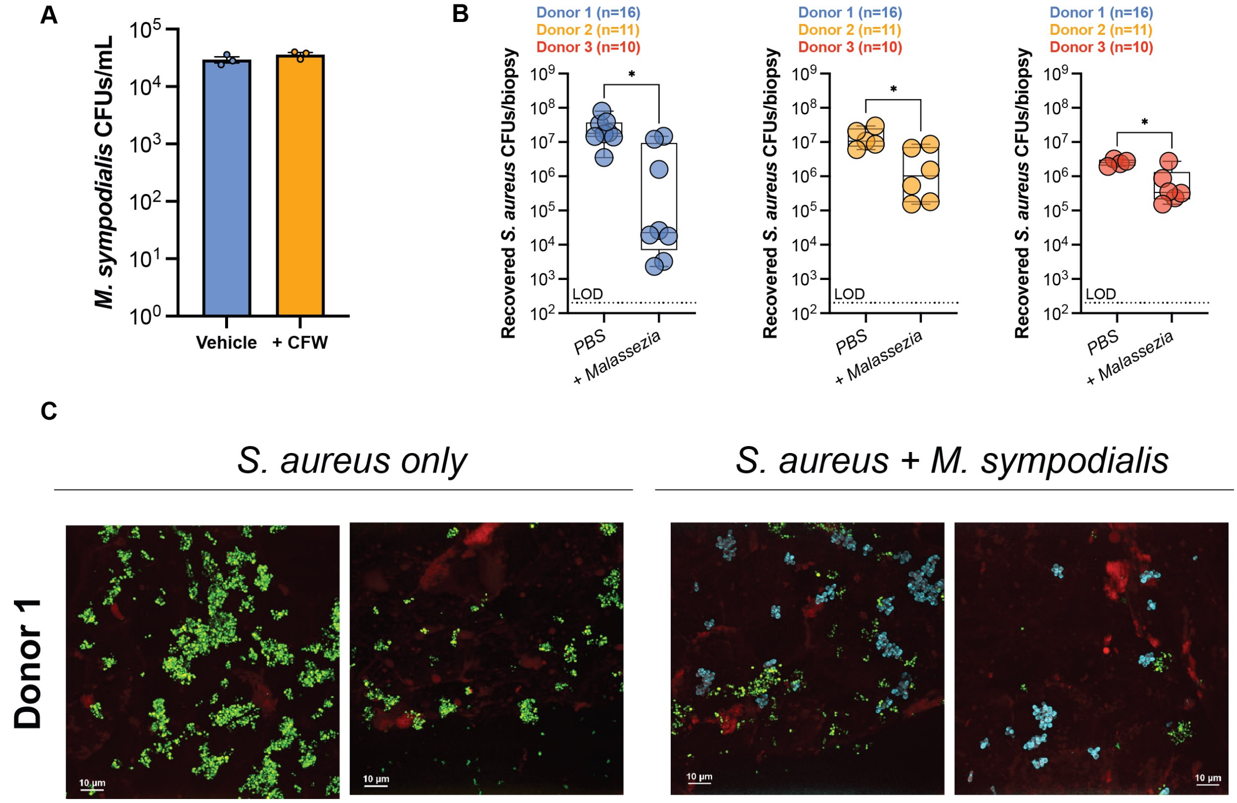


**Figure S2. Supporting data for** ***Malassezia*** **and** ***S. aureus*** **co-culture of human skin biopsies. A.** *M. sympodialis* CFUs/mL to measure viability after fluorescent labeling with the cell wall dye calcofluor white (CFW) (n=3). **B.** The data from Figure 1E separated into individual graphs for each of the three donors: donor 1 (n=16), donor 2 (n=11), donor 3 (n=10). Unpaired parametric t-test, *: p<0.05. **C.** Representative max projection images from the skin surface (red) at day 7 with *S. aureus* (green) and/or *M. sympodialis* stained with CFW (cyan). Scale bar is 10 µm. Biopsies shown are from donor 1.

B and C here are mixed up

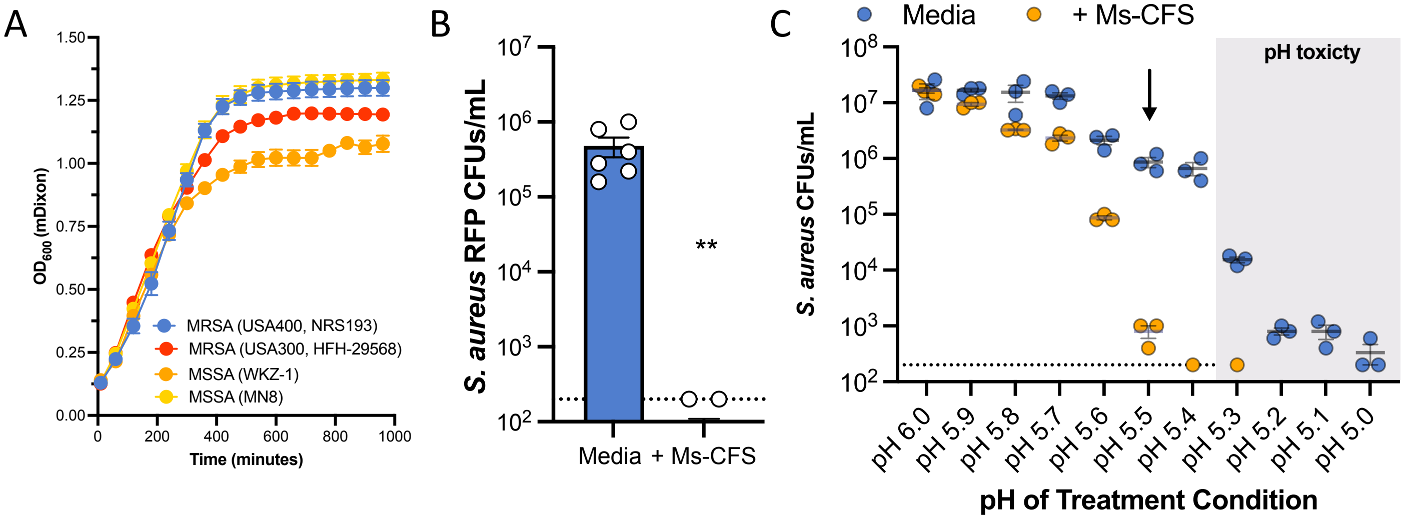


**Figure S3. *S. aureus* growth in mDixon and pH-dependent sensitivity to mDixon CFS from *M. sympodialis*. A.** Four strains of *S. aureus* cultured for 1000 min. in mDixon media at pH 6. These strains correspond to those treated in Figure 2B. The WT strain use in this study is shown blue circles (NRS193). Additional strain data in Table S1. **B.** *S. aureus* expressing red fluorescent protein (RFP) used for the experiment in Figure 2D treated with 50% Ms-CFS for 2-h compared to exposure to the pH-matched control. Unpaired t-test, **: p<0.01. Dashed line is limit of detection (200 CFUs/mL). **C.** *S. aureus* NRS193 treated for 2-h with 50% Ms-CFS (orange) or media control (blue) adjust to various pH. The grey box indicates where the pH results in media toxicity to *S. aureus*. The arrow indicates the pH (5.5) used for experiment.

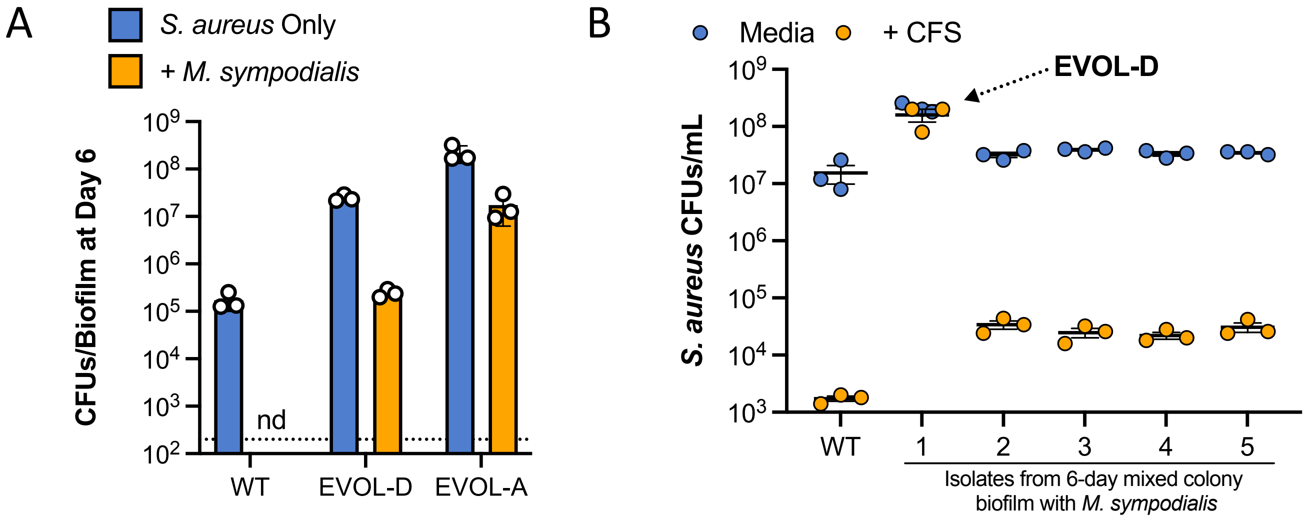


**Figure S4. Isolation of spontaneous tolerant isolate EVOL-D**. **A**. *S. aureus* CFUs/biofilm after 6 days. Each point is an independent biofilm. nd: none detected, dashed line is limit of detection (200 CFUs/biofilm). **B**. *S. aureus* CFUs/mL after 2-h exposure to media or Ms-CFS for the NRS193 WT or isolates A-E from a 6-day mixed colony biofilm with *M. sympodialis*. Each point is a technical replicate, only isolate A, EVOL-D, was cryopreserved and DNA sequenced.

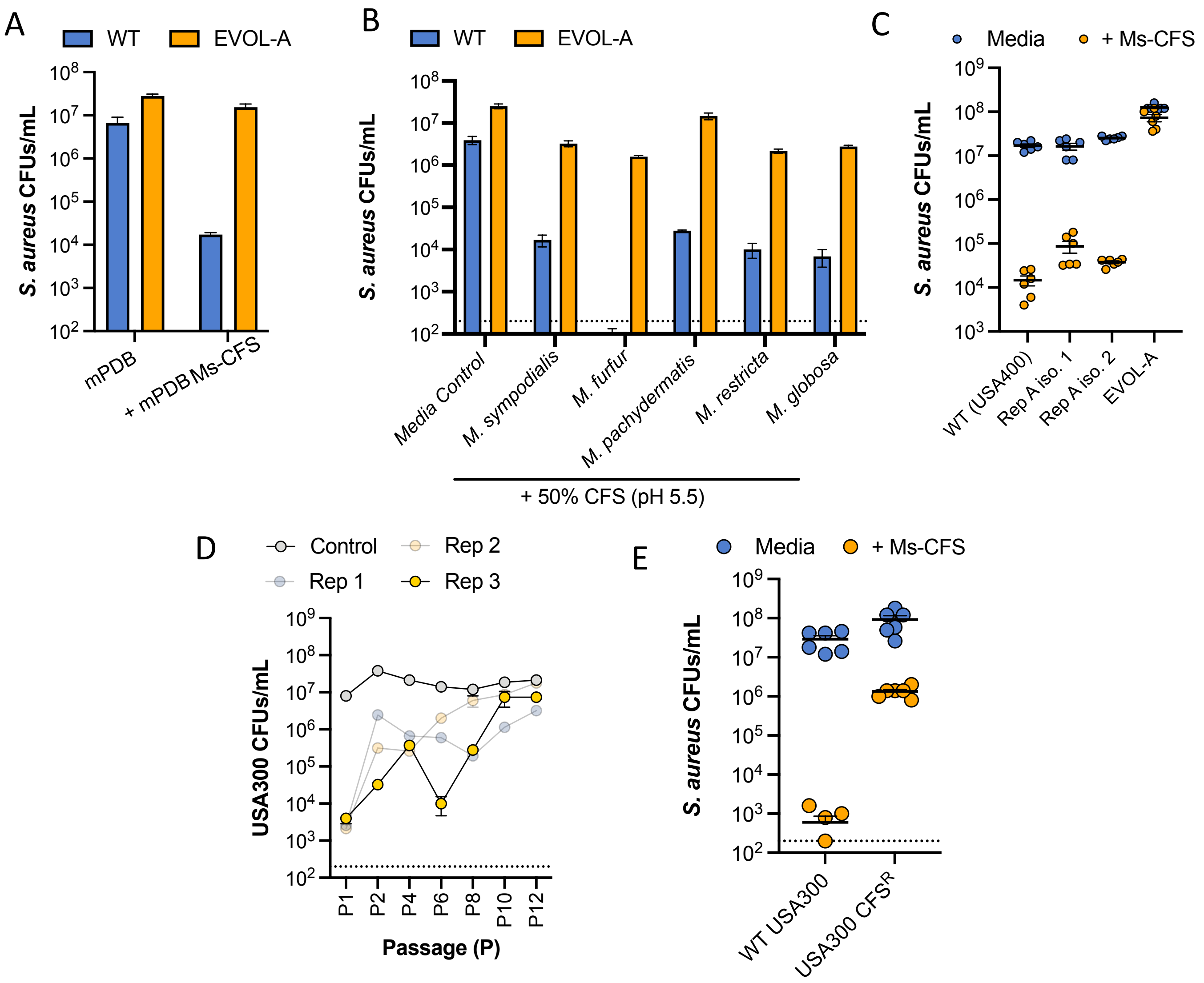


**Figure S5. Features of the isolates with evolved tolerance to *M. sympodialis* CFS. A.** *S. aureus* NRS193 WT (blue) and EVOL-A (orange) CFUs after 2-h exposure to mPDB 50% Ms-CFS or pH-matched control. **B.** *S. aureus* NRS193 WT (blue) and EVOL-A (orange) CFUs after 2-h exposure to 50% CFS from various *Malassezia* species grown in mDixon and adjusted to pH 5.5. **C.** *S. aureus* CFUs/mL after 2-h exposure to 50% *M. sympodialis* CFS. Rep A iso. 1 and Rep A iso. 2 are isolates from passage 12 of Rep A (the same as EVOL-A) that have the WT allele of *rel* (n=6). **D.** *S. aureus* USA300 CFUs/mL recovered from sequential passages in mDixon (Control) or *M. sympodialis* 50% CFS (Rep 1, Rep 2, Rep 3). Error bars (SEM) represent technical variation as the viable counts in the passaged population were measured only between each passage (P). **E.** *S. aureus* CFUs/mL of the USA300 WT or an isolate evolved from USA300 from Rep 3 P12 ($CFS^R$) after 2-h exposure to *M. sympodialis* 50% CFS (n=6). Dashed line is limit of detection (200 CFUs/mL).

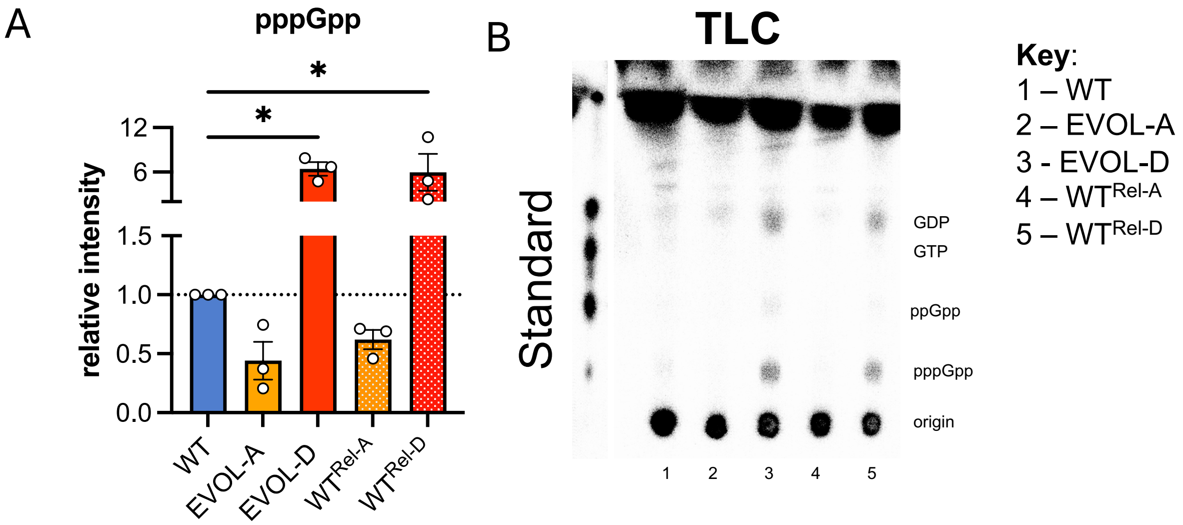


**Figure S6. pppGpp measurements in *S. aureus*. A**. Quantification of pppGpp levels relative to the NRS193 WT after induction with mupirocin. Data from three independent experiments. One-way ANOVA with Dunnett multiple comparisons test. *: $p<0.05$, error bars indicate standard deviation around the mean. **B**. Representative thin layer chromatography (TLC) plate quantifying pppGpp.

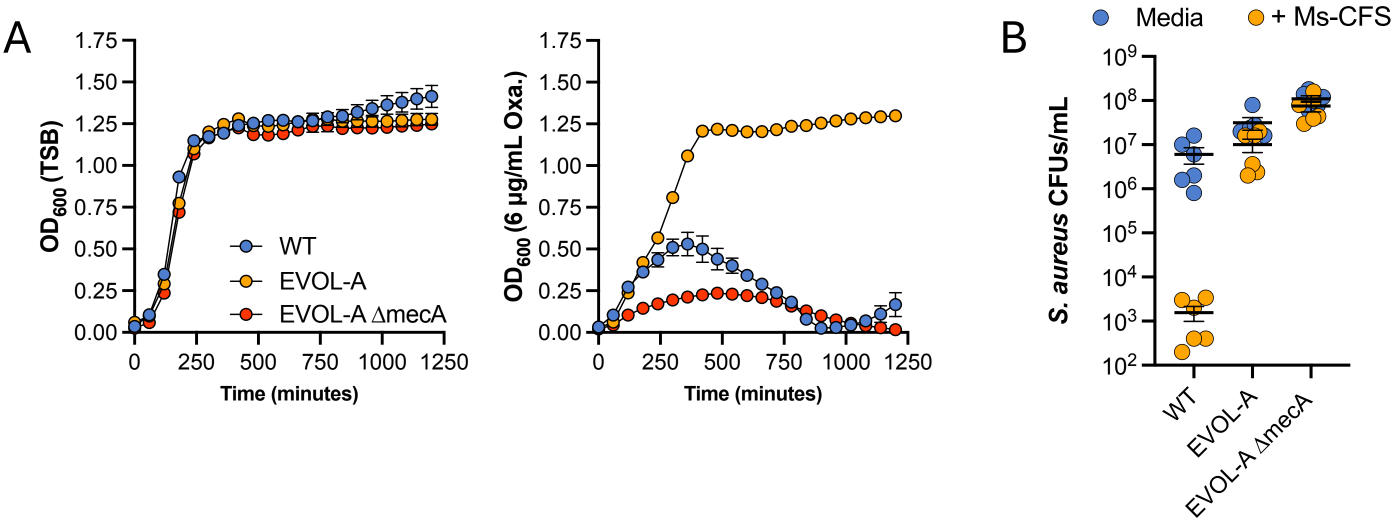


**Figure S7. Loss of *mecA* in EVOL-A**. A. Growth curve of $OD_{600}$ measurements over 20-h in TSB or TSB + 6 µg/mL oxacillin (n=4). **B**. *S. aureus* CFUs/mL after 2-h exposure to media control or Ms-CFS for NRS193 WT, EVOL-A, and the deletion of *mecA* in EVOL-A (EVOL-A Δ*mecA*). Data pooled from two independent experiments.

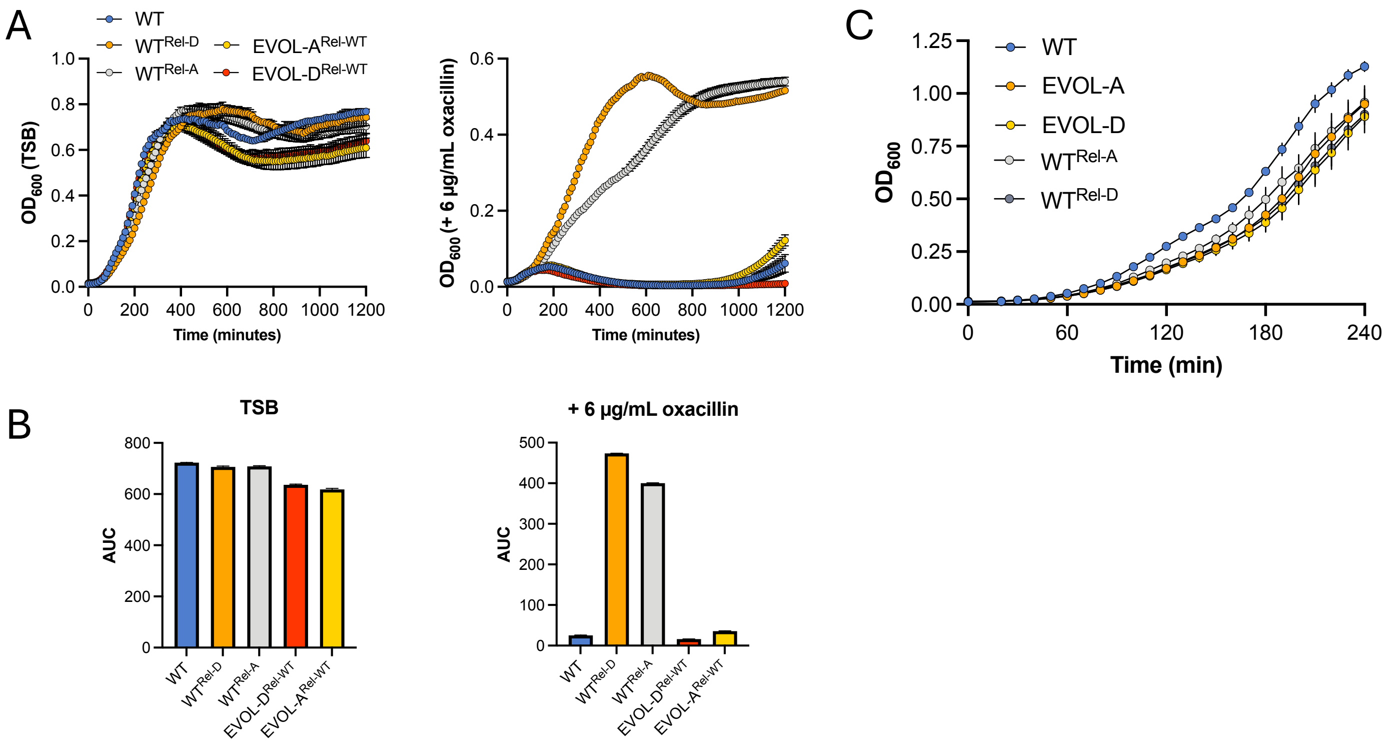


**Figure S8. Growth dynamics of Rel allele swaps in TSB with and without oxacillin.** Representative growth curves in TSB or TBS + 6 μg/mL oxacillin for 20-h with area under the curve (AUC) shown in **B. C.** Representative growth curve of first 4-h of growth in TSB with measurements at $OD_{600}$ taken at 10-min. intervals.

| # | Species | Strain | Notes | Acquisition | Link/Reference |
|---|---|---|---|---|---|
| 1 | *Staphylococcus aureus* | NRS193 (WT) | CA-MRSA, USA400, closely related to MW2 | BEI Resources, NIAID, NIH | https://www.beiresources.org/Catalog/Bacteria/NR-45992.aspx |
| 2 | *Staphylococcus aureus* | Je2 | MRSA, USA300 | BEI Resources, NIAID, NIH | https://www.beiresources.org/Catalog/Bacteria/NR-46543.aspx |
| 3 | *Staphylococcus aureus* | HFH-29568 | MRSA, USA300 | BEI Resources, NIAID, NIH | https://www.beiresources.org/Catalog/bacteria/NR-10186.aspx |
| 4 | *Staphylococcus aureus* | HFH-30364 | MRSA, USA400 | BEI Resources, NIAID, NIH | https://www.beiresources.org/Catalog/bacteria/NR-10189.aspx |
| 5 | *Staphylococcus aureus* | SR2609 | USA300, MRSA | BEI Resources, NIAID, NIH | https://www.beiresources.org/Catalog/Bacteria/NR-50507.aspx |
| 6 | *Staphylococcus aureus* | RN4850 | MSSA | BEI Resources, NIAID, NIH | https://www.beiresources.org/Catalog/bacteria/NR-45955.aspx |
| 7 | *Staphylococcus aureus* | WKZ-1 | MSSA | BEI Resources, NIAID, NIH | https://www.beiresources.org/Catalog/bacteria/NR-28984.aspx |
| 8 | *Staphylococcus aureus* | MN8 | MSSA | BEI Resources, NIAID, NIH | https://www.beiresources.org/Catalog/bacteria/NR-45918.aspx |
| 9 | *Staphylococcus aureus* | BR-VRSA | VRSA | BEI Resources, NIAID, NIH | https://www.beiresources.org/Catalog/Bacteria/NR-49120.aspx |
| 10 | *Staphylococcus aureus* | IL (Isolate F) | VISA | BEI Resources, NIAID, NIH | https://www.beiresources.org/Catalog/Bacteria/NR-45905.aspx |
| 11 | *Staphylococcus aureus* | LIM2 | VISA | BEI Resources, NIAID, NIH | https://www.beiresources.org/Catalog/bacteria/NR-45881.aspx |
| 12 | *Escherichia coli* | Nissle | | Gifted by K. Guillemin | |
| 13 | *Staphylococcus hominis* | SK119 | | BEI Resources, NIAID, NIH | https://www.beiresources.org/Catalog/Bacteria/HM-119.aspx |
| 14 | *Staphylococcus hominis* | VCU122 | | BEI Resources, NIAID, NIH | https://www.beiresources.org/Catalog/Bacteria/NR-46399.aspx |
| 15 | *Staphylococcus capitis* | SK14 | | BEI Resources, NIAID, NIH | https://www.beiresources.org/Catalog/Bacteria/HM-117.aspx |
| 16 | *Staphylococcus capitis* | VCU116 | | BEI Resources, NIAID, NIH | https://www.beiresources.org/Catalog/Bacteria/NR-46394.aspx |
| 17 | *Staphylococcus epidermidis* | NIHLM001 | | BEI Resources, NIAID, NIH | https://www.beiresources.org/Catalog/Bacteria/HM-896.aspx |
| 18 | *Staphylococcus epidermidis* | NIHLM040 | | BEI Resources, NIAID, NIH | https://www.beiresources.org/Catalog/bacteria/HM-912.aspx |
| 19 | *Staphylococcus epidermidis* | NIHLM015 | | BEI Resources, NIAID, NIH | https://www.beiresources.org/Catalog/bacteria/HM-901.aspx |
| 20 | *Staphylococcus epidermidis* | NIHLM020 | | BEI Resources, NIAID, NIH | https://www.beiresources.org/Catalog/bacteria/HM-904.aspx |
| 21 | *Staphylococcus epidermidis* | SK135 | | BEI Resources, NIAID, NIH | https://www.beiresources.org/Catalog/Bacteria/HM-118.aspx |
| 22 | *Staphylococcus epidermidis* | W23144 | | BEI Resources, NIAID, NIH | https://www.beiresources.org/Catalog/bacteria/HM-142.aspx |
| 23 | *Staphylococcus aureus* | EVOL-A | Evolved in M. sympodialis CFS, Replicate Populations A, NRS193 background | This Study | |
| 24 | *Staphylococcus aureus* | EVOL-B | Evolved in M. sympodialis CFS, Replicate Populations A, NRS193 background | This Study | |
| 25 | *Staphylococcus aureus* | EVOL-C | Evolved in M. sympodialis CFS, Replicate Populations A, NRS193 background | This Study | |
| 26 | *Staphylococcus aureus* | EVOL-D | Isolated from 6-day co-culture with M. sympoidalis, NRS193 background | This Study | |
| 27 | *Staphylococcus aureus* | EVOL-A ΔmecA | markerless deletion of mecA coding sequence | This Study | |
| 28 | *Staphylococcus aureus* | $WT^{Rel\text{-}A}$ | markerless allele swap of WT rel with the rel allele from EVOL-A; Q672* | This Study | |
| 29 | *Staphylococcus aureus* | $WT^{Rel\text{-}D}$ | markerless allele swap of WT rel with the rel allele from EVOL-D; L127V | This Study | |
| 30 | *Staphylococcus aureus* | $EVOL\text{-}A^{Rel\text{-}WT}$ | markerless allele swap of EVOL-A rel with the WT rel allele | This Study | |
| 31 | *Staphylococcus aureus* | $EVOL\text{-}D^{Rel\text{-}WT}$ | markerless allele swap of EVOL-D rel with the WT rel allele | This Study | |
| 32 | *Staphylococcus aureus* | $WT^{F128Y}$ | markerless allele swap of WT rel with the natural F128Y allele | This Study | |
| 33 | *Staphylococcus aureus* | $WT^{A567*}$ | markerless allele swap of WT rel with the natural A567* allele | This Study | |
| 34 | *Staphylococcus aureus* | $WT^{E657*}$ | markerless allele swap of WT rel with the natural E657* allele | This Study | |
| 35 | *Staphylococcus aureus* | $WT^{\Delta 11bp,658*}$ | markerless allele swap of WT rel with the natural Δ11bp allele that truncates at position 658 | This Study | |
| 36 | *Staphylococcus aureus* | S. aureusGFP | NRS193 expressing pGFP-pH plasmid, $Chloramphenicol^R$ | This Study | |
| 37 | *Staphylococcus aureus* | S. aureusRFP | RN4220 expressing pSRFPS1, $Trimethoprim^R$ | BEI Resources, NIAID, NIH | https://www.beiresources.org/Catalog/BEIPlasmidVectors/NR-51164.aspx |
| 38 | *Escherichia coli* | K-12, DC10B | Universal host for constructing plasmids for introduction into staphylococci, $streptomycin^R$ | BEI Resources, NIAID, NIH | https://www.beiresources.org/Catalog/Bacteria/NR-49804.aspx |
| 39 | *Staphylococcus aureus* | $USA300\ CFS^R$ | Evolved in M. sympodialis CFS, HFH-29568 background | This Study | . |
| 40 | *Malassezia sympodialis* | ATCC42132 | | ATCC | https://www.atcc.org/products/42132 |
| 41 | *Malassezia sympodialis* | KS013 | isolated from a healthy control | Gifted by J. Heitman | Gioti et al. 2013 |
| 42 | *Malassezia sympodialis* | KS014 | isolated from a healthy control | Gifted by J. Heitman | Gioti et al. 2013 |
| 43 | *Malassezia sympodialis* | KS269 | isolated from a patient with moderate to severe atopic eczema | Gifted by J. Heitman | Gioti et al. 2013 |
| 44 | *Malassezia sympodialis* | KS270 | isolated from a patient with moderate to severe atopic eczema | Gifted by J. Heitman | Gioti et al. 2013 |

### Table S1. Strain used in this study

| Strain | Tolerance Phenotype | Coding Mutations | | |
|---|---|---|---|---|
| EVOL-A | CFS Tolerant | rel, Q672* (CAA→TAA) | MW0270, (G)7→9, 461/684 nt | sspA, (GTTAGGGTTATCAGGATT)2→4 |
| EVOL-B | CFS Tolerant | rel, Δ7 bp (2033-2039/2190 nt) | MW0270, (G)7→9, 461/684 nt | |
| EVOL-C | CFS Tolerant | rel, C627F (TGC→TTC) | MW0270, (G)7→9, 461/684 nt | |
| EVOL-D | CFS Tolerant | rel, L127V (TTA→GTA) | MW0270, (G)7→9, 461/684 nt | |
| Rep A isolate 1 | CFS Sensitive | MW1553, T201S (ACC→AGC) | MW0270, (G)7→9, 461/684 nt | |
| Rep A isolate 2 | CFS Sensitive | MW1553, T201S (ACC→AGC) | MW0270, (G)7→9, 461/684 nt | MW1918, N8N (AAC→AAT) |
| USA300 $CFS^R$ | CFS Tolerant | rel, Q519* (CAG→TAG) | | |

**Table S2. Variant Calling in Evolved *S. aureus* strains.** Gene IDs from MW2 (GenBank: BA000033.2), except USA300 $CFS^R$

| Sequence ID | sequence # | rel mutation | notes/meta data |
|---|---|---|---|
| CAA3828468.1 | 1 | K477* | T192_T03_C01, throat isolate |
| KFA43512.1 | 1 | K484fs (495 AA) | Brady-SAP149 |
| SCT08151.1 | 1 | E511fs (514 AA) | GKP136-19, bulk milk |
| SUJ65441.1 | 1 | N524fs (528 AA) | NCTC12035 |
| **CAC6026400.1** | **1** | **A567*** | **NRS49, human/disease isolate** |
| WP_088171638 | 1 | N593* | SCPM-O-B-7906, food/clinical specimens |
| CAC9064778.1 | 1 | I637* | MOS437, human/disease isolate |
| CAC9381391.1 | 1 | A655fs (663 AA) | CF32, human/disease isolate |
| **EVS74071.1** | **2** | **E657*** | **M1493, M1518, MRSA Surveillance II** |
| SPZ99995.1 | 1 | P663fs (668 AA) | NCTC7878, feline |
| EHT36149.1 | 1 | D689fs (700 AA) | CIG1769, bacteremia isolate |
| **EUV05891.1** | **1** | **Q660fs (664 AA)** | **M0188, MRSA Surveillance II** |
| EVC10233.1 | 1 | L678fs (679 AA) | M0623, MRSA Surveillance II |

**Table S3. Identification of natural Rel C-terminal truncation variants.** Protein sequences with large deletions elsewhere in the protein were excluded. Bold indicates alleles utilized in this study.

| Primer ID | Target/purpose | Primer Sequence |
|---|---|---|
| Rel_swap_P1 | S. aureus DNA, clone Rel into pIMAY | AAAAGCGGCCGCatgaacaacgaatatccatatagtgcagatg |
| Rel_swap_P2 | S. aureus DNA, clone Rel into pIMAY | AAAACCCGGGtcattgaattttgccgtcctgactttc |
| F128Y_P1 | Q5 site directed mutagenesis of Rel | AATCGCAATAtATAACTTGCGATGATTTTC |
| F128Y_P2 | Q5 site directed mutagenesis of Rel | GCCAAAGATGTACGCGTA |
| E657_P1 | Q5 site directed mutagenesis of Rel | GCAGTTACCTaTAAATCAACCTG |
| E657_P2 | Q5 site directed mutagenesis of Rel | GTATGACCGAAATGGCTTG |
| A567_P1 | Q5 site directed mutagenesis of Rel | ACGAAATCATTGCCTATTAAA |
| A567_P2 | Q5 site directed mutagenesis of Rel | AACTTCTTGTTATTCATTTAAAGCACG |
| fs658_P1 | Q5 site directed mutagenesis of Rel | ATATTTTTGAGTTGCGTCTTTTG |
| fs658_P2 | Q5 site directed mutagenesis of Rel | AGAGGTAACTGCGTATGAC |
| mecA_P1 | Generation of ΔmecA in EVOL-A | ggtattactggaccgaatggactag |
| mecA_P2 | Generation of ΔmecA in EVOL-A | gtcttatataaggagtatattgatgcaaaacagtgaagcaatccgtaacg atg |
| mecA_P3 | Generation of ΔmecA in EVOL-A | catcaatatactccttatataagactacatttgtagtatattac |
| mecA_P4 | Generation of ΔmecA in EVOL-A | gacttgccattaattctgcttgctctacaac |
| mecA_P5 | Generation of ΔmecA in EVOL-A | gttgtgcttgttaaacttgcagaag |
| mecA_P6 | Generation of ΔmecA in EVOL-A | gctttagcactgtctcgtttagaccaatc |
| mecA_IntF | Generation of ΔmecA in EVOL-A | cagagttaatgggaccaacataacctaatag |
| mecA_IntR | Generation of ΔmecA in EVOL-A | agtagaaatgactgaacgtccg |
| PSarAP1_P1 | sarA promoter, screening insertions in pTH3 | cccagaaatacaatcactgtgtctaatgaataatttg |
| GFPph_P1 | pHlourin GFP screening primer | GTGTGTTATTCCAGCAGCTGTCAC |

| Plasmid ID | Purpose | Abx | Source |
|---|---|---|---|
| pIMAY | allelic replacement in S. aureus | chloramphenicol (E. coli and S. aureus) | Addgene (#68939), Monk et al. 2012 |
| pTH3 | shuttle plasmid for sarA P1-YFP expression | ampicillin (E. coli), chloramphenicol (S. aureus) | Addgene (#84453), de Jong et al. 2017 |
| pGFP-pH | PsarA-P1-pHlourin, GFP plasmid also used to assess changes in intracellular pH | ampicillin (E. coli), chloramphenicol (S. aureus) | This study, built by replacing YFP (GPVenus) in pTH3 with pHlourin |
| pRel-swap | Rel locus in pIMAY for Q5 site directed mutagenesis | chloramphenicol (E. coli and S. aureus) | This study |

**Table S4. Primers and Plasmids used in this study.**
